# Simulating the effect of ankle plantarflexion and inversion-eversion exoskeleton torques on center of mass kinematics during walking

**DOI:** 10.1101/2022.11.07.515398

**Authors:** Nicholas A. Bianco, Steven H. Collins, Karen Liu, Scott L. Delp

**Affiliations:** Department of Mechanical Engineering, Stanford University, Stanford, California, United States of America; Department of Computer Science, Stanford University, Stanford, California, United States of America; Department of Bioengineering, Stanford University, Stanford, California, United States of America; Department of Orthopaedic Surgery, Stanford University, Stanford, California, United States of America

## Abstract

Walking balance is central to independent mobility, and falls due to loss of balance are a leading cause of death for people 65 years of age and older. Bipedal gait is typically unstable, but healthy humans use corrective torques to counteract perturbations and stabilize gait. Exoskeleton assistance could benefit people with neuromuscular deficits by providing stabilizing torques at lower-limb joints to replace lost muscle strength and sensorimotor control. However, it is unclear how applied exoskeleton torques translate to changes in walking kinematics. This study used musculoskeletal simulation to investigate how exoskeleton torques applied to the ankle and subtalar joints alter center of mass kinematics during walking. We first created muscle-driven walking simulations using OpenSim Moco by tracking experimental kinematics and ground reaction forces recorded from five healthy adults. We then used forward integration to simulate the effect of exoskeleton torques applied to the ankle and subtalar joints while keeping muscle excitations fixed based on our previous tracking simulation results. Exoskeleton torque lasted for 15% of the gait cycle and was applied between foot-flat and toe-off during the stance phase, and changes in center of mass kinematics were recorded when the torque application ended. We found that changes in center of mass kinematics were dependent on both the type and timing of exoskeleton torques. Plantarflexion torques produced upward and backward changes in velocity of the center of mass in mid-stance and upward and forward velocity changes near toe-off. Eversion and inversion torques primarily produced lateral and medial changes in velocity in mid-stance, respectively. Intrinsic muscle properties reduced kinematic changes from exoskeleton torques. Our results provide mappings between ankle plantarflexion and inversion-eversion torques and changes in center of mass kinematics which can inform designers building exoskeletons aimed at stabilizing balance during walking. Our simulations and software are freely available and allow researchers to explore the effects of applied torques on balance and gait.

## Introduction

Walking balance is central to human health and mobility. Balance ability is especially critical to older adults who face significant health problems and injuries when mobility is impaired [1]. In particular, falls lead to over 800,000 hospitalizations per year [2] and represent a leading cause of death due to injury for persons 65 and older [1, 3]. Humans avoid falling by using corrective torques to counteract internal (e.g., motor and sensory noise) and external (e.g., uneven surfaces) perturbations [4]. However, fall risk increases when factors including muscle weakness [5, 6] and neurological impairments such as Parkinson’s disease [7–9] compromise the ability for an individual to avoid undesired center of mass motions. For example, falling after tripping is associated with reduced push-off strength [10, 11] and slower response times after a trip [12, 13] in older adults.

Exoskeleton devices may help prevent undesired center of mass motions by providing the corrective torques that individuals with muscle weakness or sensorimotor impairments are unable to produce. Ankle exoskeletons could also be used to replicate the effect of ankle plantarflexor muscles on changes in medio-lateral ground reaction forces during stance [14]. Furthermore, ankle inversion-eversion exoskeleton torques could be used to mimic the medio-lateral ankle strategy observed in humans, where the center of pressure position under the foot is shifted in response to a perturbation [15–17]. Recent studies have investigated how ankle exoskeletons affect measures related to gait stability including gait variability [18], margin of stability [19], and muscular effort [20]. Other studies have used pelvis perturbations to observe how humans counteract induced center of mass velocity changes using stepping strategies and by modulating joint torques and muscle activity [15, 16, 21–23]. For example, Vlutters et al. (2018) found that humans increase ankle plantarflexion torque in response to anterior pelvis perturbations [22]. These studies suggest that ankle and subtalar exoskeleton torques would have strong effects on fore-aft and medio-lateral walking motions, respectively. However, the current literature lacks a comprehensive understanding of how the timing of exoskeleton torques applied to the ankle joint, subtalar joint, or both joints combined lead to three-dimensional changes in center of mass and center of pressure kinematics.

Previous studies have used simulation to study walking balance using a variety of approaches [14, 24–38]. Simplified biped models have been used to study how modulating push-off [31, 35] and joint speed [25] can help stabilize walking. Kim et al. (2017) found that ankle push-off torque modulation had a strong effect on the maximum tolerable medio-lateral perturbation during walking [31]. Several studies have simulated how humans maintain walking balance using models of muscle-reflex through feedback control [29, 30, 36, 37] based on the model of Geyer and Herr (2010) [39], and others have incorporated feed-forward control strategies to model the contribution of central pattern generators to walking balance [24, 26, 33]. Three-dimensional musculoskeletal simulations have revealed the contributions of muscles to medio-lateral ground reaction forces [14, 27] and foot placement strategies after perturbation [32, 38]. John et al. (2012) used simulation to investigate how intrinsic muscle properties (i.e., force-length, force-velocity, and tendon stiffness properties) reduce kinematic changes after force perturbations applied to the pelvis, highlighting the importance of including the effects of muscle properties when studying walking balance [28]. However, no simulation study has provided mappings between exoskeleton torques applied to the ankle and subtalar joints and changes in center of mass motions, which would be valuable information for designers building exoskeletons to aid walking balance.

The goal of this study was to understand the effect of ankle plantarflexion and inversion-eversion exoskeleton torques on center of mass kinematics and center of pressure positions during unperturbed walking. To this end, we first created realistic muscle-driven simulations of walking using optimal control that track experimental gait data. We then modeled exoskeleton devices using massless actuators that applied plantarflexion and inversion-eversion torques to the ankle and subtalar joints and used forward integration to simulate their effect on walking kinematics. We analyzed these simulations to achieve two goals. First, we sought to determine how changes in center of mass kinematics and center of pressure positions differed based on different exoskeleton devices and different timings of exoskeleton torque during the gait cycle. Second, we sought to determine how intrinsic muscle properties alter center of mass and center of pressure changes induced by exoskeleton torques.

## Methods

We utilized a two-step approach to simulate the effect of ankle plantarflexion and inversion-eversion exoskeleton torques during walking (Fig. 1). We first created muscle-driven simulations of walking by tracking experimental motion capture and ground reaction force data. Massless actuators were then added to the ankle and/or subtalar joints to model the exoskeleton devices which applied torques in plantarflexion, inversion, eversion, plantarflexion and inversion, or plantarflexion and eversion. The effects of the exoskeleton torques were simulated using forward integration in OpenSim [40], where either the muscle excitations from the tracking simulation controlled the muscles in the model (“muscle-driven” simulations), or the moments generated by the muscles in the tracking simulation were applied to the model using torque actuators (“torque-driven” simulations). Exoskeleton torques were applied during the stance phase between foot-flat and toe-off, and the initial kinematic state of the model was prescribed using the state of the model from the tracking solution at the onset of exoskeleton torque. To evaluate the effect of the exoskeleton torques, we computed changes in center of mass kinematics and center of pressure positions from the forward simulations. Finally, the effect of excluding intrinsic muscle properties was evaluated by comparing the muscle-driven and torque-driven simulation results.

**Fig 1.**
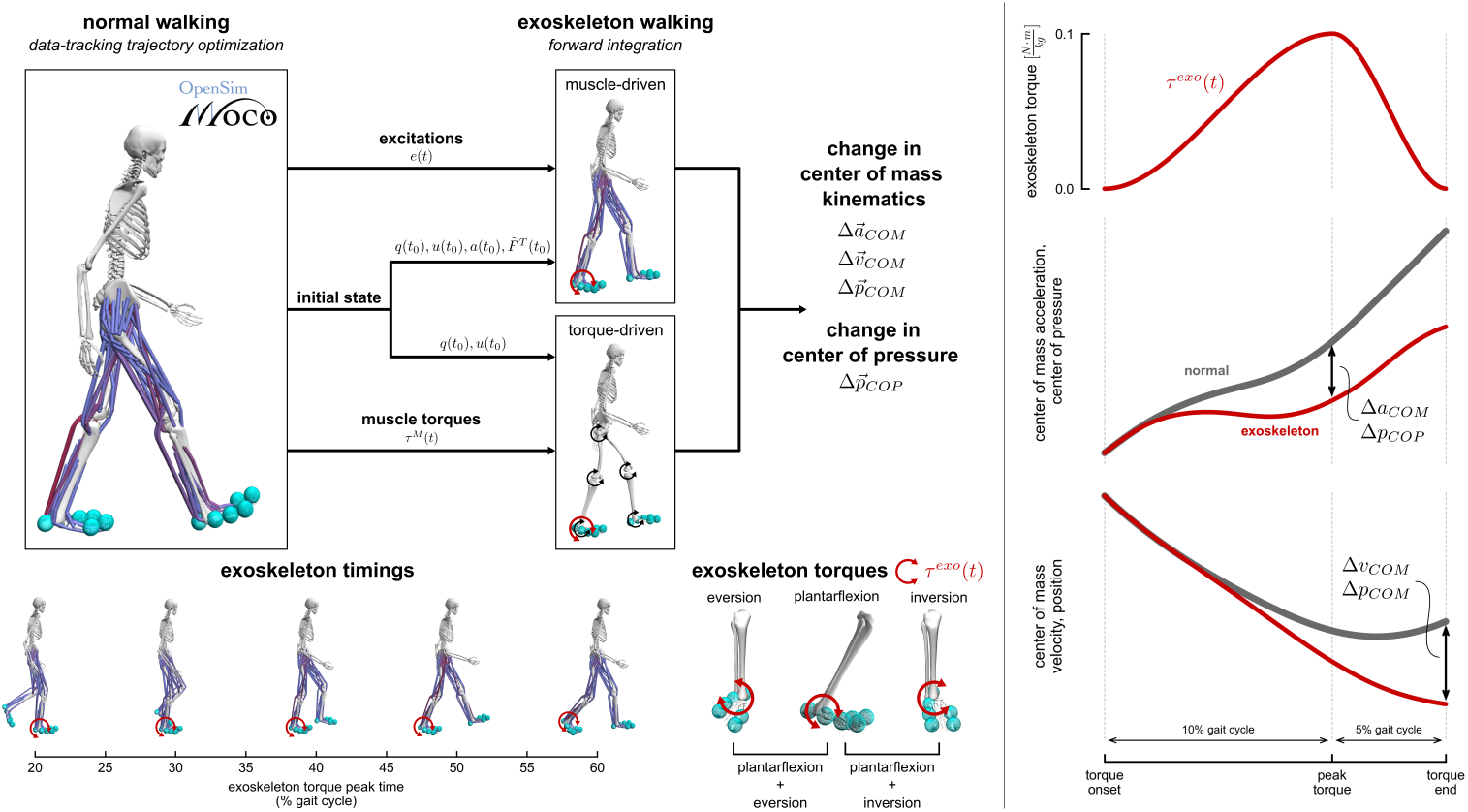
Overview of simulation approach. Left: We used a two-step approach to simulate the effect of ankle plantarflexion and inversion-eversion exoskeleton torques during walking. In the first step, we used OpenSim Moco to create muscle-driven simulations of walking that tracked experimental motion capture data. In the second step, we applied different ankle plantarflexion and inversion-eversion torques at times between 20% (foot-flat) and 60% (toe-off) of the gait cycle and simulated the effects using forward integration. We simulated five different exoskeleton torques: plantarflexion, eversion, inversion, plantarflexion plus eversion, and plantarflexion plus inversion. The initial state of each exoskeleton simulation was set using the tracking simulation state at the beginning of the applied torque. In the muscle-driven simulations, the excitations from the tracking simulation were prescribed to the muscles and torque actuators in the model. In the torque-driven simulations, the torques generated by muscles in the tracking simulation were applied to the model. Right: The exoskeleton torques had a magnitude of 0.1 N-m/kg and rise and fall times equal to 10% and 5% of the gait cycle, respectively. We computed center of mass acceleration and center of pressure changes at the time of peak exoskeleton torque, and center of mass velocity and position changes were computed at the time that exoskeleton torque ended. The curves illustrate when each quantity was calculated, but do not represent actual trajectories for each quantity.

### Experimental data

We used a previously-collected dataset from 5 healthy individuals walking on a treadmill (mean ± s.d.: age: 29.2 ± 6.3 years, height: 1.80 ± 0.03 m, mass: 72.4 ± 5.7 kg) [41]. Subjects in this previous study provided informed consent to a protocol approved by the Stanford Institutional Review Board. The data included marker trajectories, ground reaction forces, and electromyography (EMG) signals from 11 muscles in the right leg. The EMG signals were normalized based on the highest value in the filtered EMG curves for each muscle across all walking and running trials in the full dataset used in Arnold et al. (2013). For each subject, we chose a single representative gait cycle of walking at 1.25 m s^-1^ to create our tracking simulations.

### Musculoskeletal model

We used a generic musculoskeletal model with 31 degrees-of-freedom and 80 lower-extremity muscles and a torque-actuated upper-extremity developed for gait simulation [42]. The model included recent modifications to muscle passive force parameters and hip adductor-abductor muscle paths [43]. Tendon compliance was enabled only for the soleus and gastrocnemius muscles using a tendon strain of 10% at maximum isometric force based on Arnold et al. (2013) [41]. We used the muscle model developed by De Groote et al. (2016), where normalized tendon force and activation were the state variables [44]. Foot-ground contact was modeled using six smoothed Hunt-Crossley force elements applied to each foot. The contact sphere size, configuration, and parameter values were based on the generic foot-ground contact models from Falisse et al. (2019) [45]. The generic model was scaled to match each subject’s static marker data using AddBiomechanics [46]. Joint kinematics were computed from the marker data of each subject from inverse kinematics using OpenSim [40].

Passive force elements were included in our model to represent contributions from ligaments and soft tissue structures. Torsional spring forces were added to the lumbar extension (1 Nm rad^-1^ kg^-1^), bending (1.5 Nm rad^-1^ kg^-1^), and rotation (0.5 Nm rad^-1^ kg^-1^) coordinates based on experimental measurements of bending moments applied to the lumbar joint [47]. Since ankle and subtalar kinematics were important to the results of this study, we included a previously developed model of passive ankle structures using three-dimensional torsional bushing forces based on cadaver experiments that span both the ankle and subtalar joints [48]. Damping forces were applied to the ankle with coefficient 2 Nm s rad^-1^ based on experimental ankle impedance measurements [49]. Based on Falisse et al. (2022), we adjusted the orientation of the metatarsophalangeal joint axis to be perpendicular to the sagittal plane and applied a linear rotational spring force with a stiffness of 25.0 Nm rad^-1^ [50]. Damping forces with a 2 Nm s rad^-1^ coefficient were also applied to the lumbar, subtalar, and metatarsophalangeal joints, since the lumbar joint was not muscle-actuated and the subtalar and metatarsophalangeal joints did not track any experimental data (see “Normal walking simulations” below). For the remaining joints in the model, we applied passive rotational exponential spring and linear damping forces to represent ligament forces [51].

### Normal walking simulations

We created muscle-driven walking simulations that tracked the experimental data using optimal control. The optimal control problems were solved using direct collocation in OpenSim Moco [52]. We used a multi-objective cost function that both minimized muscle excitations and torque actuator effort and tracking errors between model and experimental kinematics and kinetics. We minimized errors between experimental ground reaction forces and the forces produced from the foot-ground contact model.

Model coordinate values and speeds tracked coordinate trajectories computed from inverse kinematics. Since our experimental data were collected from treadmill walking, the inverse kinematics trajectories were converted to represent overground walking data by translating the fore-aft position of the pelvis forward according to the experimental treadmill speed.

In addition to tracking ground reaction forces and joint kinematics, we also tracked body kinematics of the torso and the feet to produce more realistic walking kinematics. We tracked experimental torso body orientations and angular velocities computed from our inverse kinematics results. This approach avoided any tracking errors in the pelvis coordinates from being propagated to the torso which would occur if the lumbar joint kinematics were tracked directly. Similarly, we tracked calcaneus body orientations and angular velocities to avoid abnormal kinematics produced by the foot-ground contact model or errors propagated by hip, knee, and ankle tracking errors. This approach helped produce realistic kinematics for the subtalar and metatarsophalangeal joints which did not track any experimental data.

We imposed constraints on the optimal control problem to improve performance and avoid undesired solutions. Since our experimental gait data were approximately periodic, all model states (coordinate values, coordinate speeds, activations, and normalized tendon forces) and controls (muscle and torque excitations) were constrained to be periodic across the gait cycle. This also avoided unconstrained state variables from taking on large values at the beginning of the trajectory. We prevented the arms from intersecting with the torso and the feet from intersecting with each other by imposing a minimum distance between pairs of the respective bodies via path constraints added to the problem. Finally, the average walking speed over the gait cycle was constrained to match the experimental walking speed.

The tracking problems were solved with CasADi [53]. We used a Hermite-Simpson collocation scheme with mesh intervals at every 10 ms in the gait cycle. Each problem was solved using a constraint tolerance of 1e-4 and a convergence tolerance 1e-2; these tolerances produced good agreement between model and experimental kinematics and between muscle activity and experimental EMG data (see “Validation approach” and “Validation results”). Finally, since we needed to perform forward integration to produce our exoskeleton walking simulations, the tracking problems were solved using explicit dynamics for both skeletal and muscle dynamics. For more information about how the optimal control problem was implemented in OpenSim Moco, see S1 Appendix.

### Exoskeleton walking simulations

We simulated five different exoskeleton devices that applied one or two torques to the ankle and/or subtalar joints. The first device applied a plantarflexion torque to the ankle joint, where equal and opposite body torques were applied to the tibia and the talus. Two exoskeleton devices applied torque in eversion and inversion to the subtalar joint, where equal and opposite body torques were applied to the talus and calcaneus. The last two exoskeleton devices applied torque to the ankle and subtalar joints simultaneously (i.e.,“plantarflexion plus eversion” and “plantarflexion plus inversion”). Each exoskeleton device was modeled using massless torque actuators with a a rise time equal to 10% of the gait cycle and a fall time equal to 5% of the gait cycle based on a common four parameter control design used in previous exoskeleton experimental studies [54]. We chose a peak torque of 0.1 Nm kg-1 because it was both large enough to produce an effect on center of mass motion and small enough to be used for closed-loop control of balance in real devices without destabilizing the user. The exoskeleton device torques had peak times between 20% and 60% of the gait cycle at 5% intervals which occurred between the average times of foot-flat and toe-off of the five subjects. Each simulation began at the onset of exoskeleton torque, and the model initial state was set using the state from the normal walking simulation. The effect of each exoskeleton device was evaluated using forward simulation performed with a 4th-order Runge-Kutta-Merson time-stepping integrator (integrator accuracy: 1e-6) [55]. The forward simulations were performed when exoskeleton torques were applied; therefore, each simulation had length equal to 15% of the gait cycle.

In each exoskeleton simulation, the contributions from muscles were applied to the model using two different approaches that addressed both goals of our study. In the first approach, we applied the muscle excitations from the tracking simulation to the original model with muscle actuators included (i.e., “muscle-driven” simulations). In this approach, muscle forces could change based on muscle intrinsic properties (e.g., force-length and force-velocity relationships) as the applied exoskeleton torque produced changes in kinematics. In the second approach, we first computed the moments generated by the muscles in the normal walking simulations. We then replaced the muscles in the model with torque actuators that applied the muscle-generated moments (i.e., “torque-driven” simulations). In this approach, the joint torques applied to the model could not change in response to the exoskeleton devices. The differences between the muscle-driven simulations and the torque-driven simulations reveal the influence of the intrinsic muscle properties.

We computed changes in center of mass kinematics and right foot center of pressure positions to assess the effect of each exoskeleton device in the forward simulations.

Changes in center of mass acceleration and in center of pressure were computed at the time of peak exoskeleton torque (i.e., “exoskeleton torque peak time”), since these quantities were most related to force-level information (Fig. 1, right panel). Changes in center of mass velocity and position were computed at the end of the exoskeleton torque (i.e., “exoskeleton torque end time”) to measure the full effect of each exoskeleton on the center of mass state. Center of mass kinematic quantities were made dimensionless based on the recommendations of Hof (1996) [56]. Center of mass positions were normalized by center of mass height (*h_COM_*, 1.03 ± 0.02 m) and accelerations were normalized by gravitational acceleration (*g*). Center of mass velocities were normalized by the term 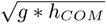 to convert to the non-dimensional Froude number. Center of pressure was normalized by an estimated foot width of approximately 11 cm based on the generic contact sphere configuration we used for all subjects.

### Validation approach

To validate the kinematics and kinetics of our simulations, we followed guidelines suggested by Hicks et al. (2015) [57]. We computed the RMS errors between experimental and model marker trajectories from inverse kinematics. We ensured that the value and timing of the joint angles computed from inverse kinematics were consistent with the previous experimental study [41]. The RMS errors between experimental and tracking simulation coordinate kinematics and ground reaction forces were computed. Lastly, we computed the RMS error between simulation muscle activations and EMG signals; we accounted for the electromechanical delay in muscle force production by applying a delay of 40 ms to the EMG signals [58]. The timing of foot-flat, heel-off, and toe-off and step width values were compared to typical healthy walking values. The kinematics of the subtalar and metatarsophalangeal joints were compared to experimental kinematics previously reported in the literature [59, 60].

### Statistical testing

We performed two sets of statistical tests to address the goals of our study. The first set of tests compared the effects of the exoskeleton devices on changes in center of mass kinematics and center of pressure positions relative to the normal walking simulations. For these tests, we employed a linear mixed model (fixed effect: exoskeleton device; random effect: subject) with 25 observations for each test (5 subjects and 5 exoskeleton devices). Individual tests were performed for each time point in the gait cycle, model actuator type (muscle-driven or torque-driven), and direction of center of mass or center of pressure change (i.e., fore-aft, vertical, or medio-lateral change). The second set of tests evaluated if the changes in center of mass kinematics and center of pressure positions from the torque-driven simulations were significantly different from the changes produced from the muscle-driven simulations. We again employed a linear mixed model (fixed effects: exoskeleton and actuator type; random effect: subject) with 50 observations for each test (5 subjects, 5 exoskeletons, and 2 actuator types).

Individual tests were performed for each time point in the gait cycle and direction of center of mass or center of pressure change. For each linear mixed model, we performed analysis of variance (ANOVA) tests and Tukey post-hoc pairwise tests with a significance level of *α* = 0.05 [61]. The statistical tests were performed with R [62–67].

## Results

### Changes in center of mass kinematics: muscle-driven simulations

Plantarflexion, eversion, and inversion torques produced changes in fore-aft and medio-lateral center of mass velocity in the muscle-driven simulations compared to the normal walking condition (Fig. 2, Fig. S7; Tukey post-hoc tests, p *<* 0.05). Our results coudl be generally grouped into two main phases of the gait cycle: mid-stance (25% to 55% of the gait cycle) and push-off (60% to 65% of the gait cycle). Exoskeleton plantarflexion torque produced backward and lateral changes in velocity during mid-stance and a slight forward change in velocity during push-off. Eversion exoskeleton torque produced lateral velocity changes during mid-stance and a medial change during push-off. Inversion exoskeleton torques produced the opposite effect: medial velocity changes during mid-stance and a lateral change during push-off. During late mid-stance, eversion and inversion torques produced backward and forward changes in center of mass velocity, respectively. Eversion and inversion torques produced larger medio-lateral velocity changes compared to plantarflexion torque, whereas plantarflexion torque produced larger fore-aft velocity changes compared to eversion and inversion torques. No significant changes in fore-aft or medio-lateral velocity were produced by any exoskeleton torque at 60% of the gait cycle. See S1 Appendix for center of mass acceleration and position results.

**Fig 2.**
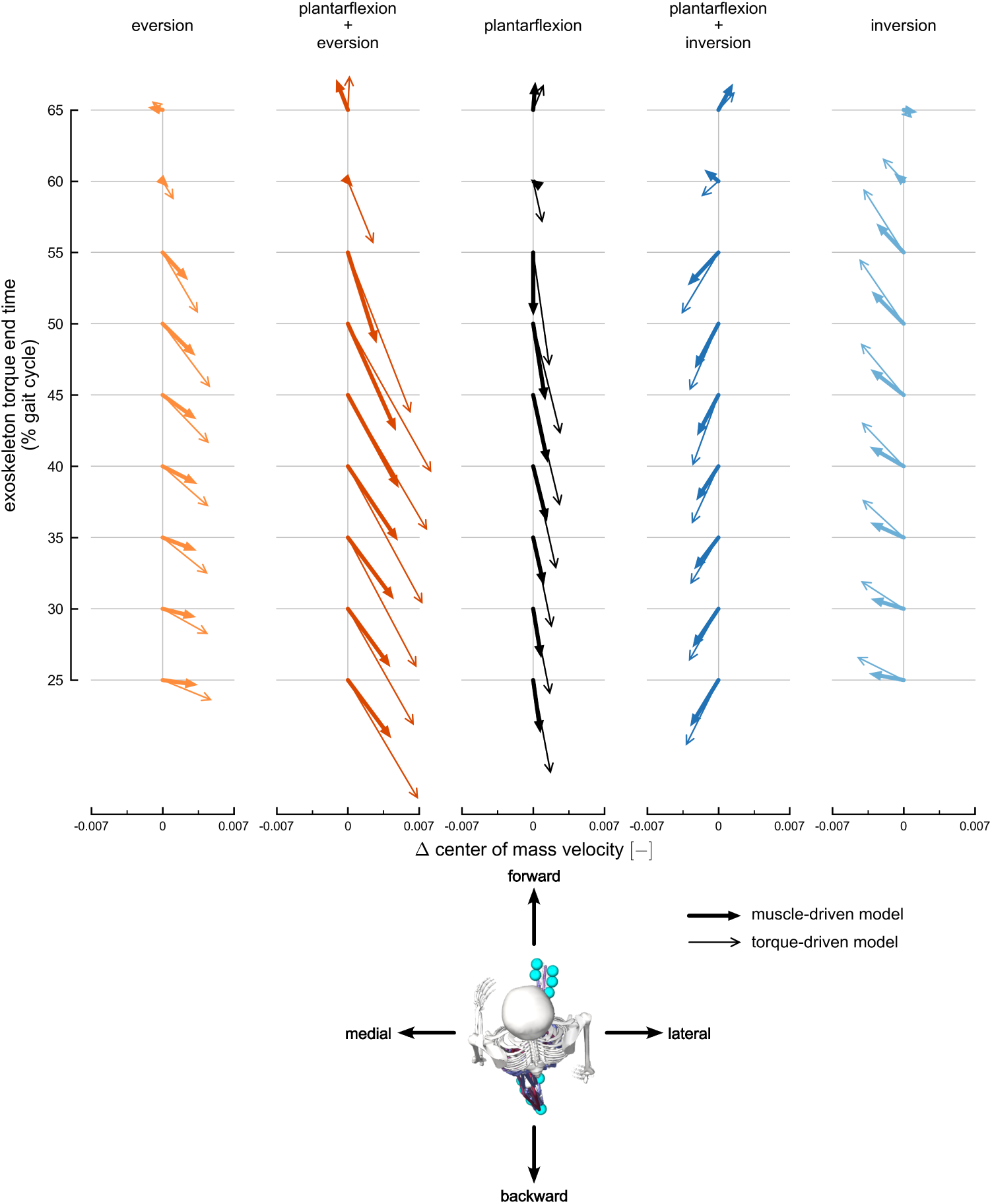
Transverse-plane changes in center of mass velocity. The change in center of mass velocity, calculated at exoskeleton torque end time, projected onto the transverse plane. Velocity changes are normalized to the dimensionless Froude number. Columns represent velocity changes for each exoskeleton torque condition: eversion (light orange), plantarflexion plus eversion (dark orange), plantarflexion (black), plantarflexion plus inversion (dark blue), and inversion (light blue). Thick arrows with filled heads represent changes using the muscle-driven model; thin arrows with open heads represent results using the torque-driven model. Each column includes velocity changes at different exoskeleton torque end times, ranging from 25% (bottom) to 65% (top) of the gait cycle. The horizontal arrow directions are medio-lateral changes in velocity, and the vertical arrow directions are fore-aft changes in velocity. The horizontal axes provide scales for medio-lateral velocity changes, and the fore-aft changes represented by each arrow are scaled to match the horizontal axes. The maximum transverse velocity change observed across both muscle-driven and torque-driven conditions was 0.056 m s^-1^.

Plantarflexion plus eversion and plantarflexion plus inversion torques produced changes in fore-aft and medio-lateral center of mass velocity that combined the velocity changes produced by individual plantarflexion, eversion, and inversion torques (Fig. 2, Fig. S7; Tukey post-hoc tests, p *<* 0.05). Lateral changes in velocity from plantarflexion plus eversion torques were larger than both the eversion and plantarflexion torques during mid-stance. Medial changes in velocity from plantarflexion plus inversion torque were smaller in magnitude than changes produced by inversion torque during mid-stance, since the plantarflexion torque component produced a lateral change in velocity. The plantarflexion plus eversion and plantarflexion plus inversion torques produced a backward change in fore-aft velocity during mid-stance and a forward change during push-off.

Exoskeleton torques also produced changes in the vertical center of mass velocity (Fig. 3, Fig. S7; Tukey post-hoc tests, p *<* 0.05). Plantarflexion torque produced upward changes in vertical center of mass velocity during early mid-stance and during late mid-stance to push-off. Plantarflexion plus eversion torques also produced upward changes in velocity during early mid-stance and during late mid-stance to push-off, whereas the plantarflexion plus inversion torques produced upward changes in velocity at all time points. Eversion and inversion torques produced no changes in vertical center of mass velocity, and no exoskeleton torque produced a downward change in velocity.

**Fig 3.**
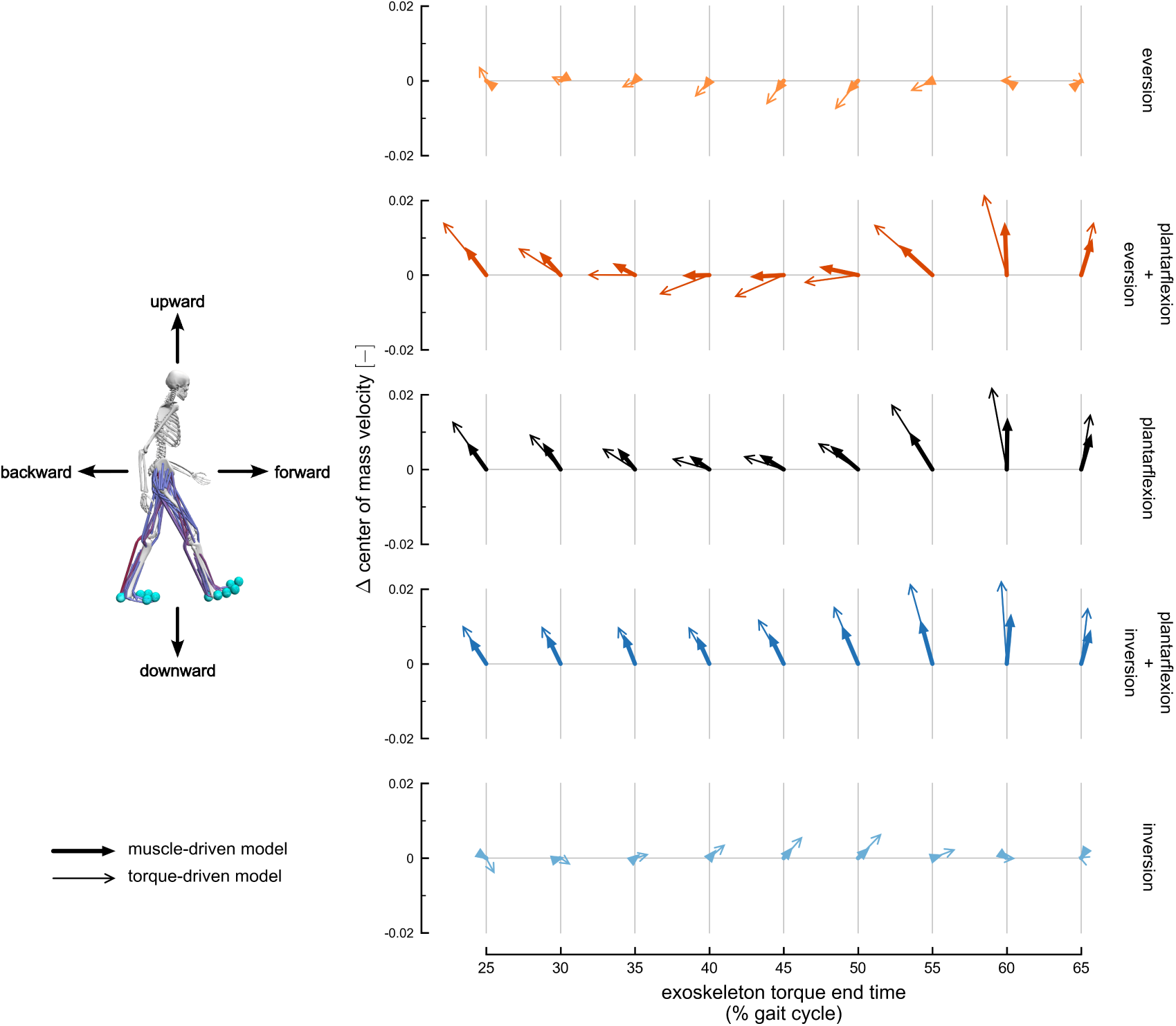
Sagittal-plane changes in center of mass velocity. The change in center of mass velocity, calculated at exoskeleton torque end time, projected onto the sagittal plane. Velocity changes are normalized to the dimensionless Froude number. The arrows represent the velocity changes averaged across subjects. Rows represent velocity changes for each exoskeleton torque condition: eversion (light orange), plantarflexion plus eversion (dark orange), plantarflexion (black), plantarflexion plus inversion (dark blue), and inversion (light blue). Thick arrows with filled heads represent changes using the muscle-driven model; thin arrows with open heads represent results using the torque-driven model. Each row includes velocity changes at different exoskeleton torque end times, ranging from 25% (left) to 65% (right) of the gait cycle. The horizontal arrow directions are fore-aft changes in velocity, and the vertical arrow directions are vertical changes in velocity. The vertical axes provide scales for vertical velocity changes, and the fore-aft changes represented by each arrow are scaled to match the vertical axis. The maximum transverse velocity change observed across both muscle-driven and torque-driven conditions was 0.072 m s^-1^.

### Changes in center of pressure positions: muscle-driven simulations

Plantarflexion, eversion, and inversion torques produced changes in fore-aft and medio-lateral right foot center of pressure positions in the muscle-driven exoskeleton simulations (Fig. 4, Fig. S9; Tukey post-hoc tests, p *<* 0.05). Plantarflexion torques shifted the center of pressure position forward and medially during mid-stance. Eversion torque shifted the center of pressure position medially, while inversion torque shifted the center of pressure laterally. During late mid-stance, eversion torque shifted the center of pressure forward and inversion torque shifted the center of pressure backward. During early mid-stance, plantarflexion torque produced larger fore-aft center of pressure changes compared to inversion-eversion torques, but changes from inversion-eversion torques exceeded changes from plantarflexion torque in late mid-stance. Finally, inversion-eversion torques produced larger medio-lateral changes in the center of pressure compared to plantarflexion torque.

**Fig 4.**
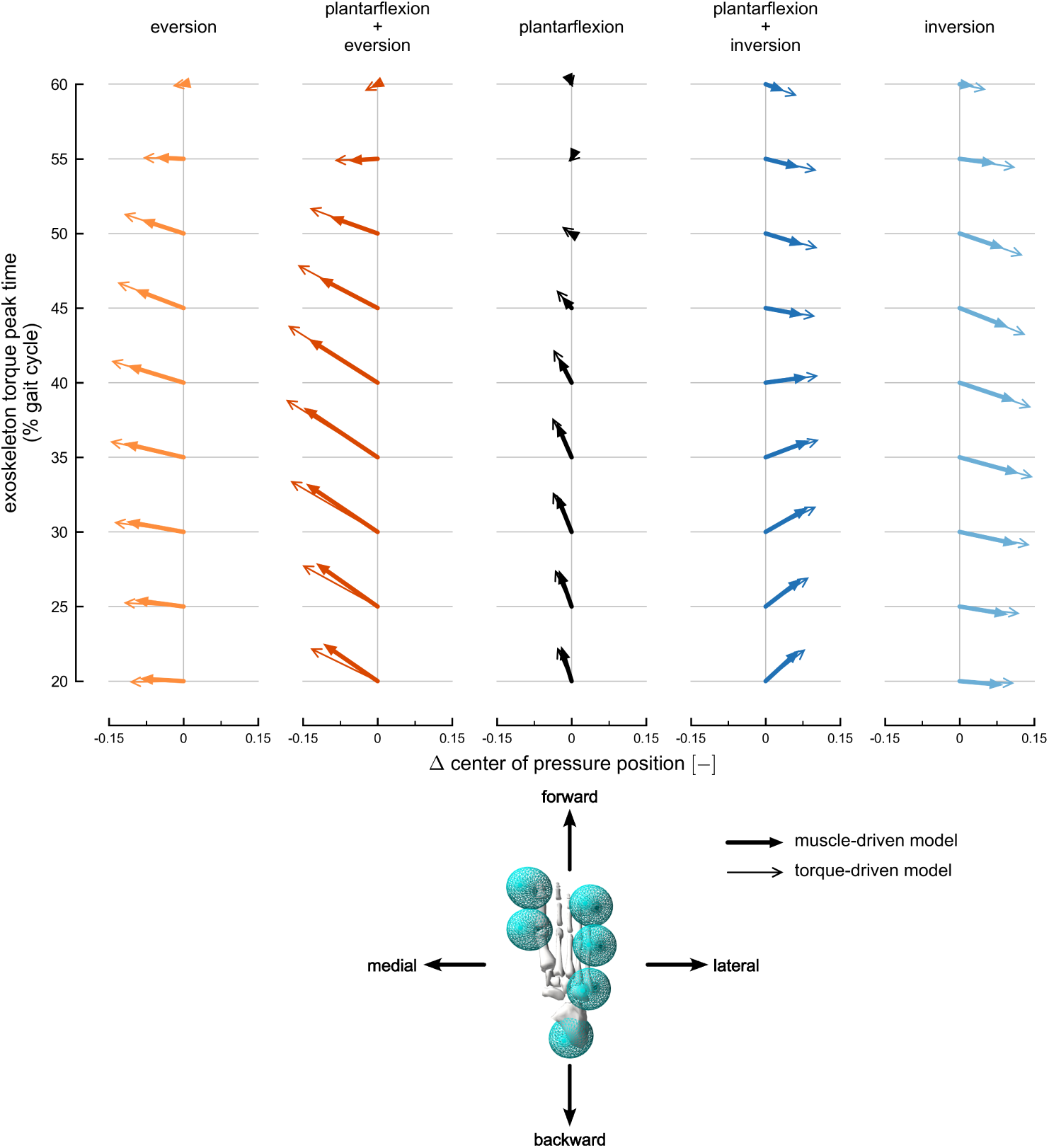
Changes in right foot center of pressure position. The change in right foot center of pressure position, calculated at exoskeleton peak time, in the sagittal plane. The arrows represent the position changes normalized by foot width and averaged across subjects. Rows represent position changes for each exoskeleton torque condition: eversion (light orange), plantarflexion plus eversion (dark orange), plantarflexion (black), plantarflexion plus inversion (dark blue), and inversion (light blue). Thick arrows with filled heads represent changes using the muscle-driven model; thin arrows with open heads represent results using the torque-driven model. Each row includes position changes at different exoskeleton timings, ranging from 20% (bottom) to 60% (top) of the gait cycle. The horizontal arrow directions are medio-lateral changes in position, and the vertical arrow directions are fore-aft changes in position. The horizontal axes provide scales for medio-lateral position changes, and the fore-aft changes represented by each arrow are scaled to match the horizontal axes. The maximum center of pressure position change observed across both muscle-driven and torque-driven conditions was 0.024 m.

Plantarflexion plus eversion and plantarflexion plus inversion exoskeleton torques produced changes in fore-aft and medio-lateral center of pressure positions that combined the changes produced by individual plantarflexion, eversion, and inversion torques (Fig. 4, Fig. S9; Tukey post-hoc tests, p *<* 0.05). In early mid-stance, the plantarflexion torque component of each exoskeleton dominated the fore-aft center of pressure changes. Combining plantarflexion and inversion torques reduced the forward shift in center of pressure positions observed with plantarflexion-only torque in mid-stance since inversion torque tended to shift the center of pressure backward.

### Torque-driven simulation results

Changes in center of mass velocity produced by the torque-driven simulations followed the same pattern as with the muscle-driven simulations, but with larger effects for many of the exoskeleton devices and timings (Fig. S7, diamonds; Tukey post-hoc tests, p *<* 0.05). The plantarflexion, plantarflexion plus eversion, and plantarflexion plus inversion torques produced significantly larger backward velocity changes during mid-stance and upward velocity changes during push-off. All exoskeletons produced significantly larger vertical velocity changes in early mid-stance and medio-lateral velocity changes in late mid-stance. Intrinsic muscle properties had a particularly large effect on center of mass kinematic changes for some exoskeleton torques and timings. For example, including muscle properties reduced changes in transverse-plane velocities from plantarflexion and plantarflexion plus eversion exoskeleton torques by nearly a factor of two during mid-stance (Fig. 2). Finally, no significant differences in center of pressure changes were detected between muscle-driven and torque-driven simulations (Fig. S9).

### Validation results

The RMS errors between experimental and model marker trajectories from inverse kinematics had a mean value of 0.7 cm and a max value of 2.8 cm across all lower-limb markers and subjects. The joint angles computed from inverse kinematics had peak values and timings consistent with those reported in the previous simulation study [41]. The mean RMS errors between experimental coordinate data and the coordinate trajectories from the tracking simulations were all less than 2 degrees, except for the left ankle which only had slightly larger errors (2.1 ± 0.8 degrees;Fig. S10). The mean RMS errors between experimental ground reaction force data and the tracking simulation forces were less than 1% bodyweight for fore-aft and medio-lateral forces and less than 5% bodyweight for vertical forces (Fig. S11). Muscle excitations generally matched EMG signals well: RMS errors were less than 0.15 for all muscles except the medial gastrocnemius and the anterior gluteus medius, which had errors of 0.2 and 0.21, respectively (Fig. S12, S1 Appendix).

The timing of foot-flat (12.5 ± 2.0% of the gait cycle) and toe-off (63.6 ± 1.4% of the gait cycle) from the tracking simulations matched well to those reported by Perry and Burnfield (2010) [68]. The timing of heel-off (39.4 ± 7.1% of the gait cycle) was later than the timing reported by Perry and Burnfield (31% of the gait cycle), but still occurred during the time range expected for the terminal stance phase (31% to 50% of the gait cycle). Step widths from the tracking simulations (0.17 ± 0.03 m) matched well to previously reported experimental values [69]. The subtalar joint, which did not track experiment data, was between -5 and 5 degrees, everted during stance, and inverted near toe-off which matched experimental kinematics [59, 60]. The metatarsophalangeal joint angle peaked at 30 degrees in extension near toe-off which matched the simulation study from Falisse et al. (2022) [50].

## Discussion

This study provides relationships between ankle and subtalar exoskeleton torques and three-dimensional changes in center of mass velocity during the stance phase of walking. We found that plantarflexion torque had a stronger effect on center of mass velocity during mid-stance compared to toe-off. In addition, plantarflexion torque reduced the forward velocity of the center of mass during mid-stance and increased forward velocity at push-off. Eversion and inversion torques primarily affected the medio-lateral velocity of the center of mass. The medio-lateral effects from subtalar torques were stronger than those from ankle plantarflexion torques, while plantarflexion torque had a stronger effect on vertical velocity compared to the subtalar torques. Plantarflexion plus eversion and plantarflexion plus inversion torques combined the kinematic changes we observed from individual exoskeleton torques. The maximum change in the magnitude of center of mass velocity we observed in the muscle-driven simulation was 0.072 m s^-1^, which is 6% of the 1.25 m s^-1^ walking speed used in our simulations. Vlutters et al. (2016) found larger velocity changes (e.g., 0.2-0.3 m s^-1^) after applying force perturbations to the pelvis [21].

During mid-stance, changes in the right foot center of pressure positions were generally in the opposite direction of the changes in center of mass kinematics. Inversion-eversion exoskeleton torques produced the largest medio-lateral changes in center of pressure which led to the medio-lateral changes in center of mass kinematics. Similarly, plantarflexion exoskeleton torque produced larger forward changes in center of pressure compared to inversion-eversion torques. Center of pressure changes were reduced as fewer contact spheres were in contact with the ground during the late mid-stance and push-off phases. The largest forward change in center of mass velocity occurred at the end of push-off when the center of pressure could no longer shift forward and produce a backwards change in center of mass velocity. In addition, ankle plantarflexion velocity peaked near toe-off (60% of the gait cycle; Fig. S10) resulting in the largest positive ankle plantarflexion exoskeleton powers we observed (Fig. S13). Finally, we found that changes in the fore-aft center of mass velocity were not strongly related to the center of mass position relative to the center of pressure (Fig. S14).

The torque-driven simulations produced greater center of mass changes compared to the muscle-driven simulations, suggesting that the effect of intrinsic muscle properties should not be excluded when using simulations to design ankle exoskeleton devices. Even when muscle excitations are not allowed to change, muscles have an important effect on how joint and center of mass kinematics evolve in response to an external force applied to the musculoskeletal system. Changes in joint angles and velocities from external forces produce changes in lengths and velocities of muscles, which lead to changes in muscle-tendon forces due to the muscle fiber force-length relationships, the fiber force-velocity relationship, and tendon elasticity. These forces tend to reduce changes in joint-level kinematics which in turn limit changes in center of mass kinematics. The results from this study are consistent with the findings reported by John et al. (2012) and provide quantitative comparisons specific to ankle devices that exoskeleton designers may find useful. For example, we found that including muscle properties could reduce the change in center of mass velocity by nearly a factor of two after applying ankle plantarflexion torque. Therefore, exoskeletons aimed at inducing kinematic changes may need torques larger than estimations from torque-driven simulations because muscles will produce forces that tend to reduce the effects of exoskeleton torques.

Some limitations of our simulation approach should be considered when interpreting our results. First, we only simulated the effect of exoskeleton torques when applied during normal walking. Therefore, it is unclear whether the relationships we observed between exoskeleton torque and center of mass state would apply to walking conditions that differ from typical gait. Second, since muscle activations were unchanged by the applied exoskeleton torques, we only simulated center of mass and center of pressure changes over a short time window (15% of the gait cycle or 165 ms on average). This time window was chosen to be a compromise between previously reported delays in muscle activity response after perturbation (100-150 ms) [16, 70] and torque lengths typically used in exoskeleton experiments [54]. Simulating for longer durations would require a model that accurately represented how the nervous system modulates muscle activity in response to exoskeleton torques, which our model did not include. Therefore, it is unknown from our results how the center of mass changes in response to exoskeleton torques over longer durations. Furthermore, a recent study has shown that exoskeleton assistance should be applied faster than the human physiological response to improve standing balance [71], so future studies should carefully consider the relationships between perturbations and the timing of assistance to optimal assist walking balance. Third, we applied the exoskeleton torques using massless actuators, so we were not able to examine effects of device mass on the center of mass changes. Finally, we did not model inter-subject variation in the orientation and kinematics of the subtalar joint axis [59, 60], which could affect changes in medio-lateral kinematics produced from exoskeleton torques.

Future studies could build upon our simulations to improve predictions of the effect of exoskeletons on walking kinematics. Models of human motor control should be incorporated in simulations to elucidate how both reflexes and voluntary neural commands drive human kinematic responses to exoskeleton torques. Simulations that train neural control models using approaches based on single-shooting [72, 73] or reinforcement learning [74] would allow for predictions of gait kinematics beyond the time range used in this study. In addition, including short range stiffness and other muscle properties not included in this study could provide more realistic muscle behavior and improve predictions of exoskeleton torques on changes in center of mass motions. Finally, we focused our analysis on the ankle joint since many experimental exoskeleton studies have found large benefits from ankle assistance (e.g., metabolic cost), and ankle end-effectors are typically easier to design compared to the knee and hip. However, exoskeleton torques applied to these joints would also lead to strong responses in the center of mass motions; our simulation framework can be easily extended to study these joints.

We chose to simulate healthy, unperturbed walking in this study since no comprehensive mapping between ankle exoskeleton torque and center of mass motion existed for any type of gait prior to this study. Future studies focusing on participants with neuromuscular deficits and other conditions that lead to minimal gait deviations from normal walking may still benefit directly from our results, and studies focusing on larger gait deviations can use these results to generate hypotheses for new assistance strategies to aid balance. Additionally, this study can be a useful reference for future work studying how to optimally assist perturbed walking, which can encompass a wide variety of motions that stem from normal walking. Finally, future studies can leverage our simulation framework to study optimal balance assistance strategies targeted at gait pathologies or perturbed walking.

Exoskeleton designers may use the results from this study as a guide for developing devices to control the center of mass during walking. We recommend applying plantarflexion exoskeleton torque during mid-stance to control the center of mass velocity primarily in the fore-aft direction. While we only simulated plantarflexion torque which decreased fore-aft velocity, dorsiflexion torque (or a reduction in an applied plantarflexion torque) could be used to produce the opposite velocity change at a specific time in the gait cycle (e.g., to increase the center of mass forward velocity during mid-stance). While previous work from simple biped models has suggested that plantarflexion torque has a strong effect on medio-lateral kinematics [31], our results suggest that eversion and inversion torques have the strongest effect on medio-lateral center of mass motions. Furthermore, inversion and eversion torques did not produce the vertical velocity changes observed with plantarflexion torque, which may be an undesired secondary effect when controlling medio-lateral balance. While we applied the same torque magnitude to the ankle and subtalar joints, different combinations of torque magnitudes at these joints could facilitate precise control over the direction of velocity changes. For example, the subtalar exoskeleton torques produced a fore-aft velocity later in mid-stance, and these changes could be negated using small plantarflexion torques to produce a purely medio-lateral velocity change. Furthermore, since the ankle and subtalar joints are adjacent, exoskeleton devices like the Anklebot from Roy et al. (2009), which can apply moments to both joints simultaneously, can approximate the plantarflexion plus inversion and plantarflexion plus eversion devices we evaluated in this study [75]. Feedback control based on the relationships between exoskeleton torques and timings and the kinematic changes we observed could be used to develop assistance strategies for improving walking stability. Finally, future studies employing simulation-based device design methods should use muscle-driven models to include the effects of intrinsic muscle properties on responses to exoskeleton torques.

## Conclusion

We used musculoskeletal simulation to evaluate the effect of ankle plantarflexion and inversion-eversion exoskeleton torques on walking kinematics. This work clarifies how exoskeleton torques applied to the ankle and subtalar joints can alter center of mass kinematics and center of pressure positions during walking. Our results showed that the exoskeleton torques we applied produced clear trends in center of mass kinematics. In addition, when we removed muscles from the exoskeleton simulations, the changes in center of mass kinematics increased, highlighting how muscle properties influence the dynamics of human-device interactions. Designers can use these results as a guide for building exoskeletons aimed at improving gait stability. We invite researchers to use our freely available simulation results (https://simtk.org/projects/balance-exo-sim) and software (https://github.com/stanfordnmbl/balance-exo-sim) to build upon our work.

## Acknowledgments

We thank Nicos Haralabidis and Jennifer Maier for helping review the manuscript.

## Supporting information

S1 Appendix: Optimal control details and additional results

### Optimal control problem

We solved optimal control problems that tracked experimental gait data to create our simulations of normal walking. Each problem was solved using direct collocation with the OpenSim Moco software package [52]. Custom problems with multiple objectives and constraints were constructed using the *MocoStudy* interface. In the following descriptions, OpenSim and Moco class names are denoted with italics.

#### Cost function

The cost function consisted of five terms with individual weights that minimized control effort and tracked experimental kinematics and kinetics (Eq. 1). Each term is integrated over one gait cycle of walking with initial time *t*_0_ and final time *t_f_* . In the equations below, the bolded symbols represent vector quantities, the tilde symbol denotes normalized quantities, and the hat symbol denotes experimental data.

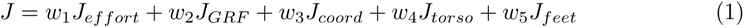

We used *MocoControlGoal* to minimize the sum of squared muscle excitations (80 muscles) and torque actuator controls (3 lumbar and 10 arm actuators), integrated over the gait cycle (Eq. 2). We divided this term by the distance traveled by the model, *d*, to encourage the model to walk forward.

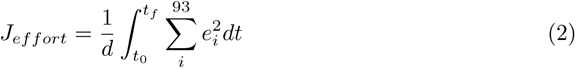

We used *MocoContactTrackingGoal* to minimize the sum of squared errors between model and experimental ground reaction forces, integrated over the gait cycle (Eq. 3). Both the model and experimental forces were normalized based on the peak force magnitudes of the experimental forces.

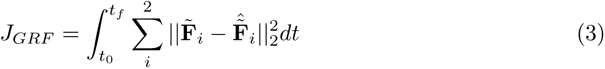

We used *MocoStateTrackingGoal* to minimize the sum of squared errors between model and experimental coordinate values, *q*, and speeds, *u*, integrated over the gait cycle (Eq. 4). The experimental coordinate values (joint angles and pelvis positions) and speeds (joint velocities and pelvis linear velocities) were the trajectories computed from the *InverseKinematicsTool* in OpenSim. The weights for individual coordinate tracking errors, *β_i_*, were normalized by the average standard deviation in each coordinate across three gait cycles. Since ankle kinematics were important to the goals of this study, we doubled the weight applied to the ankle angle tracking errors. Since our experimental data were collected from treadmill walking, the pelvis fore-aft position was translated forward according to the experimental treadmill speed to represent overground walking. The subtalar and metatarsophalangeal joints did not track experimental data since we did not have reliable experimental kinematics for these joints. We also did not track lumbar coordinates since we directly tracked torso orientations. Finally, we did not track the vertical position of the pelvis which improved ground reaction force tracking. Therefore, out of the 31 model coordinates, we tracked 25 coordinate values and 26 coordinate speeds.

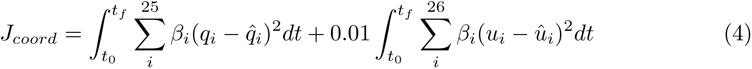

We used *MocoOrientationTrackingGoal* and *MocoAngularVelocityTrackingGoal* to track experimental torso and calcaneus body kinematics (Eq. 5-6). The experimental torso and calcaneus body kinematics were computed by first applying our inverse kinematics results to the model and computing body orientations and angular velocities. *MocoOrientationTrackingGoal* computes an angle-axis representation of the rotation matrix between the model and experimental body frames. Minimizing the angle, *θ*, from this representation minimizes the tracking error between the model and experimental body frame orientations. *MocoAngularVelocityTrackingGoal* computes the error between model and experimental three-dimensional angular velocities, ***ω***.

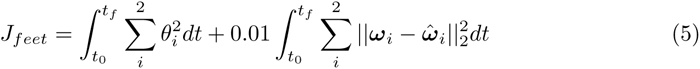

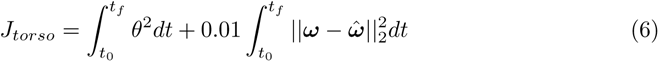

The weights for each cost function term are listed in Table A1. These weights were chosen manually such that the experimental data were tracked as closely as possible while also producing good agreement between simulated muscle activity and experimental electromyography data. We also computed the contributions of each term to the total cost function value in the final tracking solutions (Table A2).

**Table A1.** Tracking optimization cost function weights.

| weight | value |
| --- | --- |
| $w_1$ | 5 |
| $w_2$ | 7500 |
| $w_3$ | 25 |
| $w_4$ | 10 |
| $w_5$ | 10 |

#### Problem constraints

We used *MocoPeriodicityGoal* to implement endpoint constraints so that all model states and controls were periodic across the gait cycle (i.e., initial trajectory values were equal to final trajectory values). We used *MocoFrameDistanceConstraint* to prevent the model’s arms from intersecting with the torso, and to prevent the feet from intersecting with each other. Finally, we used *MocoAverageSpeedGoal* to constrain the average walking speed of the model to match the experimental walking speed.

**Table A2.** Tracking optimization cost function weights. The contributions of individual cost function terms in our tracking optimization problem to the overall cost, expressed as a percentage of the total cost function value. Results are reported as mean ± standard deviation across subjects.

| cost function term | contribution to total cost (%) |
| --- | --- |
| $J_{effort}$ | 45.0 ± 6.7 |
| $J_{GRF}$ | 15.7 ± 6.6 |
| $J_{coord}$ | 20.2 ± 6.8 |
| $J_{torso}$ | 4.6 ± 0.6 |
| $J_{feet}$ | 14.5 ± 1.8 |

#### Solver settings

We solved each problem in Moco using *MocoCasADiSolver* [53]. We used a Hermite-Simpson collocation scheme with mesh intervals at every 10 ms in the gait cycle. Each problem was solved using a constraint tolerance of 1e-4 and a convergence tolerance 1e-2 which led to good agreement between our solutions and experimental data. The tracking problems were solved using explicit dynamics for both skeletal and muscle dynamics so that the solutions could be reproduced with forward integration, which was necessary for the exoskeleton torque simulations. We used a forward difference scheme to compute function derivatives in CasADi since this reduced optimization times but did not negatively affect problem convergence. A list of important solver settings can be found in Table A3.

**Table A3.** *MocoCasADiSolver* settings. The solver settings used for each tracking optimization problem in OpenSim Moco. *t_cycle_* represents gait cycle length which was used to compute the number of mesh intervals used for each subject.

| solver setting | value |
| --- | --- |
| transcription_scheme | ‘hermite-simpson’ |
| optim_constraint_tolerance | 10<sup>−4</sup> |
| optim_convergence_tolerance | 10<sup>−2</sup> |
| num_mesh_intervals | $\frac{t_{cycle}}{0.01s}$ |
| multibody_dynamics_mode | ‘explicit’ |
| optim_finite_difference_scheme | ‘forward’ |
| scale_variables_using_bounds | true |

### Center of mass acceleration and position results

Changes in center of mass acceleration and position from the muscle-driven simulations generally reflected the results we observed for the center of mass velocity changes, with some differences (Fig. S2 to Fig. S6, Fig. S8). Changes in vertical center of mass positions were produced by all exoskeleton torques during late mid-stance. Similarly, changes in center of mass acceleration were produced by plantarflexion exoskeleton torque during mid-stance. Changes in center of mass position were observed in late mid-stance for the inversion and plantarflexion plus inversion torques. Finally, no significant changes in position changes were detected in early mid-stance from plantarflexion torque.

Changes in center of mass acceleration and position produced by the torque-driven simulations generally reflected the results we observed for the center of mass velocity changes, with some small differences in the fore-aft direction (Fig. S6 and Fig. S8, diamonds; Tukey post-hoc tests, p *<* 0.05). During late mid-stance, all torque-driven simulations produced significantly larger changes in fore-aft center of mass position and acceleration. Specifically, inversion torque produced significantly larger changes in forward center of mass acceleration during mid-stance.

### Muscle activity validation

We compared muscle activations to experimental electromyography (EMG) to validate our simulated muscle activity predictions. Individual RMS errors between muscle activations and EMG signals can be found in Table A4.

**Table A4.** RMS errors between muscle activations and electromyography. The RMS errors between simulated muscle activations and experimental EMG signals in the right leg. The EMG signals were delayed by 40 ms to account for the electromechanical delay in muscle force production [58]. Since muscle activation and EMG are both dimensionless quantities between 0 and 1, the RMS errors also lie between 0 and 1.

| muscle | RMS error |
| --- | --- |
| gluteus maximus (superior) | 0.06 |
| gluteus maximus (middle) | 0.06 |
| gluteus maximus (inferior) | 0.06 |
| gluteus medius (anterior) | 0.21 |
| gluteus medius (middle) | 0.09 |
| gluteus medius (posterior) | 0.08 |
| rectus femoris | 0.08 |
| semimembranosus | 0.09 |
| semitendinosus | 0.09 |
| vastus lateralis | 0.06 |
| vastus medialis | 0.06 |
| biceps femoris long head | 0.09 |
| lateral gastrocnemius | 0.13 |
| medial gastrocnemius | 0.20 |
| soleus | 0.11 |
| tibialis anterior | 0.11 |

**Fig S2.**
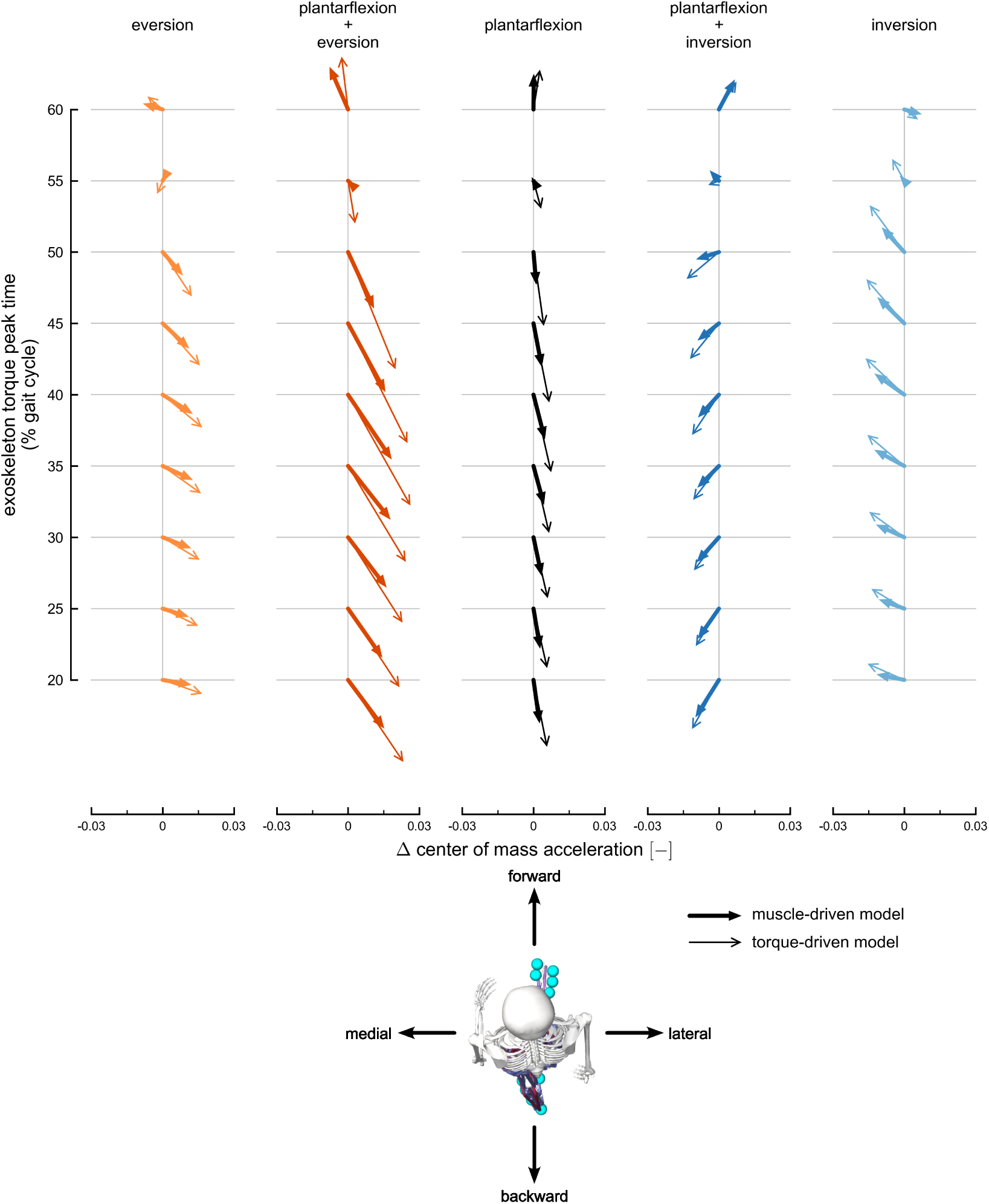
Transverse-plane changes in center of mass acceleration. The change in center of mass acceleration, calculated at exoskeleton torque peak time, projected onto the transverse plane. The arrows represent acceleration changes normalized by gravitational acceleration and averaged across subjects. Columns represent acceleration changes for each exoskeleton torque condition: eversion (light orange), plantarflexion plus eversion (dark orange), plantarflexion (black), plantarflexion plus inversion (dark blue), and inversion (light blue). Thick arrows with filled heads represent changes using the muscle-driven model; thin arrows with open heads represent results using the torque-driven model. Each column includes acceleration changes at different exoskeleton timings, ranging from 20% (bottom) to 60% (top) of the gait cycle. The horizontal arrow directions are medio-lateral changes in acceleration, and the vertical arrow directions are fore-aft changes in acceleration. The horizontal axes provide scales for medio-lateral acceleration changes, and the fore-aft changes represented by each arrow are scaled to match the horizontal axis. The maximum transverse acceleration change observed across both muscle-driven and torque-driven conditions was 0.55 m s^-2^.

**Fig S3.**
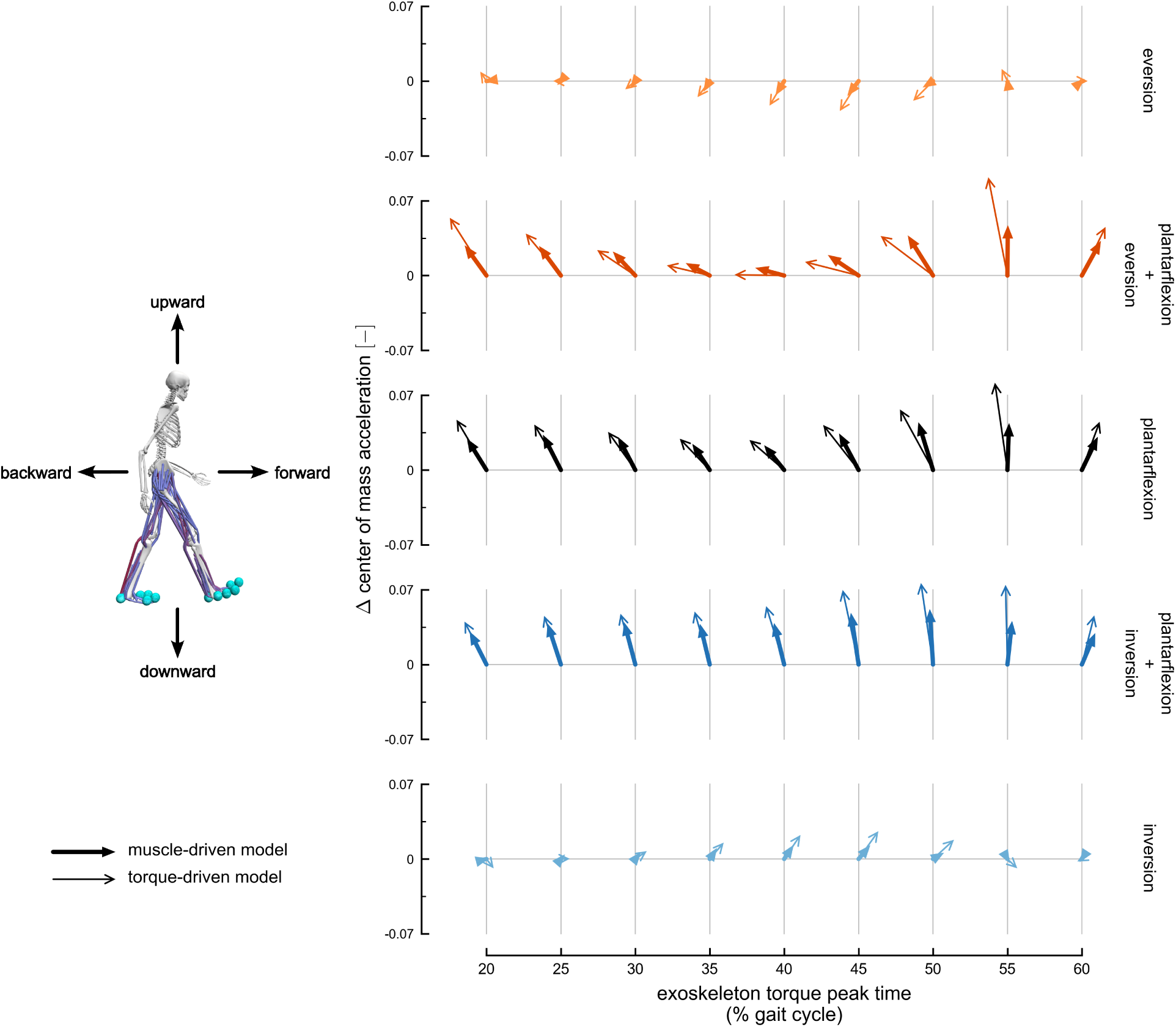
Sagittal-plane changes in center of mass acceleration. The change in center of mass acceleration, calculated at exoskeleton torque peak time, projected onto the sagittal plane. The arrows represent acceleration changes normalized by gravitational acceleration and averaged across subjects. Rows represent acceleration changes for each exoskeleton torque condition: eversion (light orange), plantarflexion plus eversion (dark orange), plantarflexion (black), plantarflexion plus inversion (dark blue), and inversion (light blue). Thick arrows with filled heads represent changes using the muscle-driven model; thin arrows with open heads represent results using the torque-driven model. Each row includes acceleration changes at different exoskeleton timings, ranging from 20% (left) to 60% (right) of the gait cycle. The horizontal arrow directions are fore-aft changes in acceleration, and the vertical arrow directions are vertical changes in acceleration. The vertical axes provide scales for vertical acceleration changes, and the fore-aft changes represented by each arrow are scaled to match the vertical axis. The maximum sagittal acceleration change observed across both muscle-driven and torque-driven conditions was 0.93 m s^-2^.

**Fig S4.**
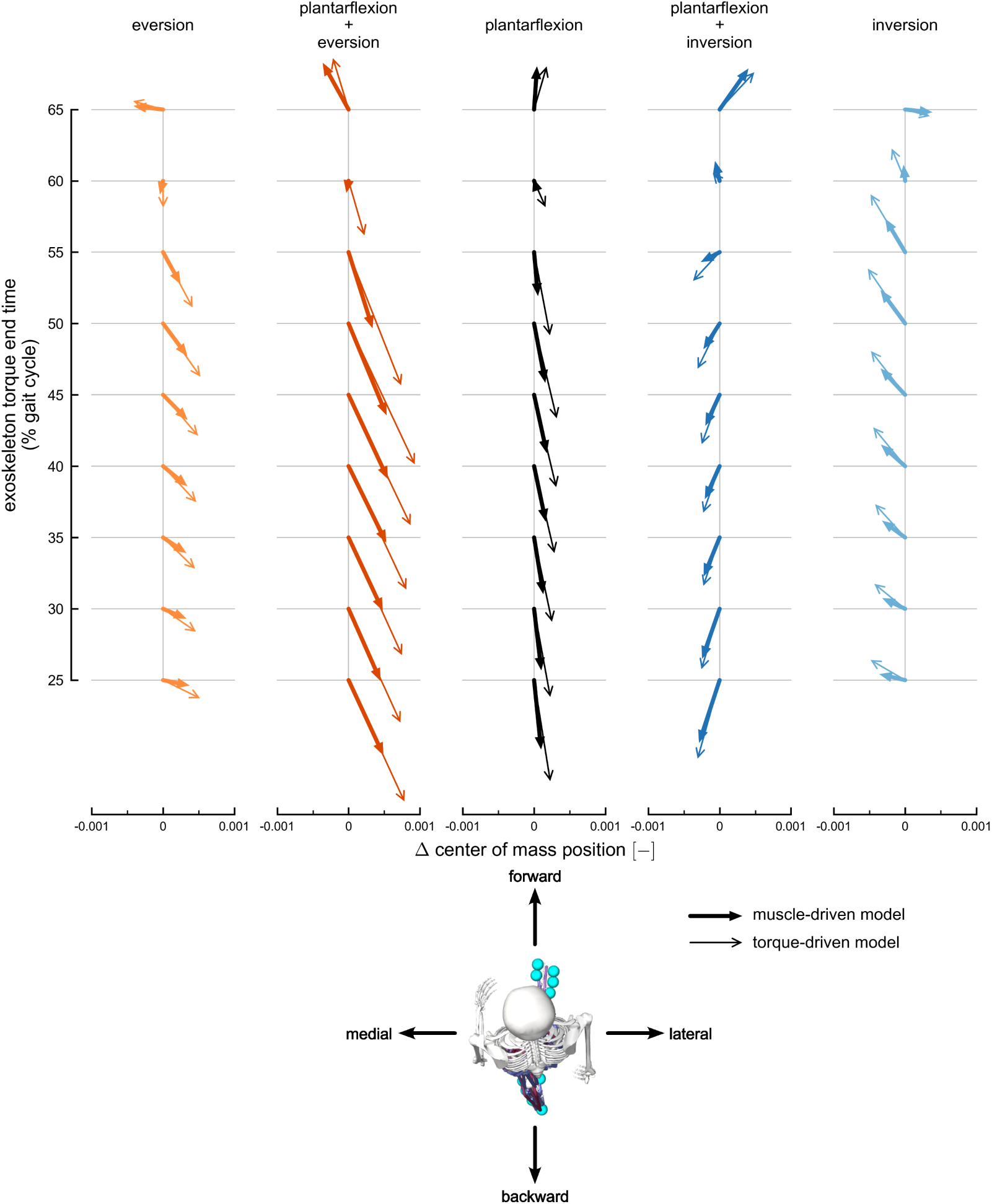
Transverse-plane changes in center of mass position. The change in center of mass position, calculated at exoskeleton torque end time, projected onto the transverse plane. The arrows represent position changes normalized by center of mass height and averaged across subjects. Columns represent position changes for each exoskeleton torque condition: eversion (light orange), plantarflexion plus eversion (dark orange), plantarflexion (black), plantarflexion plus inversion (dark blue), and inversion (light blue). Thick arrows with filled heads represent changes using the muscle-driven model; thin arrows with open heads represent results using the torque-driven model. Each column includes position changes at different exoskeleton timings, ranging from 25% (bottom) to 65% (top) of the gait cycle. The horizontal arrow directions are medio-lateral changes in position, and the vertical arrow directions are fore-aft changes in position. The horizontal axes provide scales for medio-lateral position changes, and the vertical changes represented by each arrow are scaled to match the horizontal axes. The maximum transverse position change observed across both muscle-driven and torque-driven conditions was 0.002 m.

**Fig S5.**
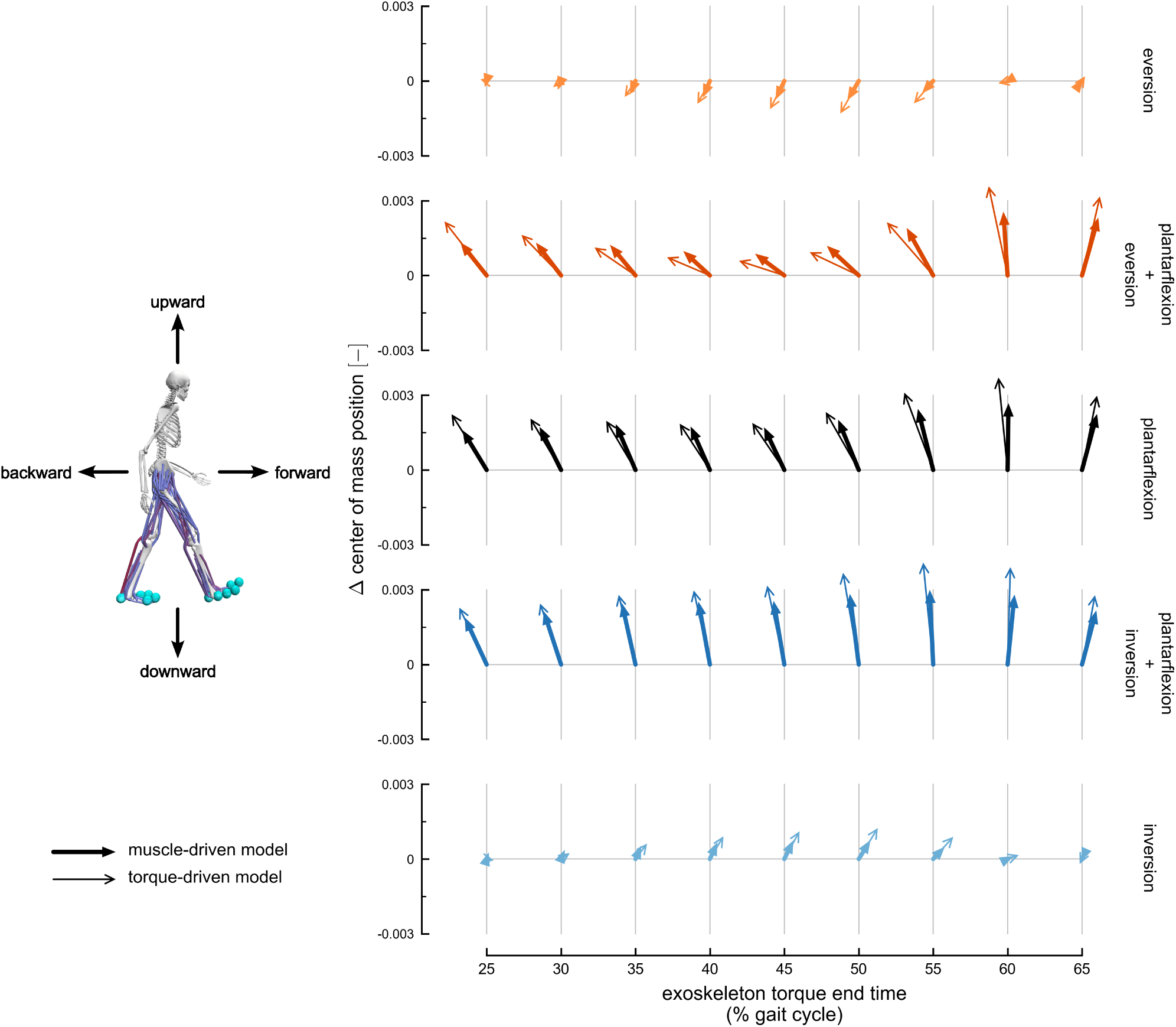
Sagittal-plane changes in center of mass position. The change in center of mass position, calculated at exoskeleton torque end time, projected onto the sagittal plane. The arrows represent position changes normalized by center of mass height and averaged across subjects. Rows represent position changes for each exoskeleton torque condition: eversion (light orange), plantarflexion plus eversion (dark orange), plantarflexion (black), plantarflexion plus inversion (dark blue), and inversion (light blue). Thick arrows with filled heads represent changes using the muscle-driven model; thin arrows with open heads represent results using the torque-driven model. Each row includes position changes at different exoskeleton timings, ranging from 25% (left) to 65% (right) of the gait cycle. The horizontal arrow directions are fore-aft changes in position, and the vertical arrow directions are vertical changes in position. The vertical axes provide scales for vertical position changes, and the fore-aft changes represented by each arrow are scaled to match the vertical axis. The maximum sagittal position change observed across both muscle-driven and torque-driven conditions was 0.004 m.

**Fig S6.**
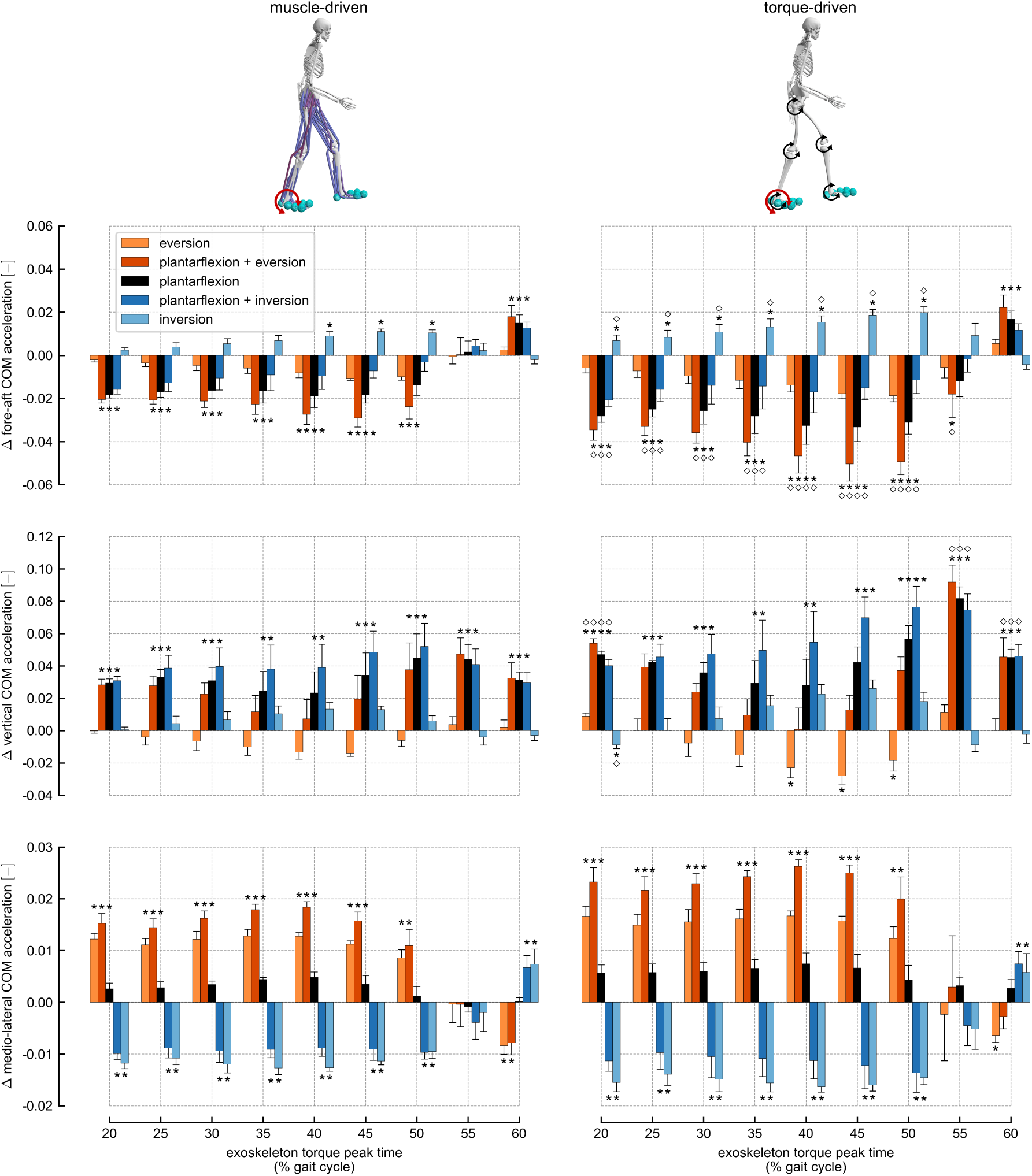
Changes in center of mass acceleration. The change in center of mass acceleration, calculated at exoskeleton torque peak time, for each exoskeleton torque condition: eversion (light orange), plantarflexion plus eversion (dark orange), plantarflexion (black), plantarflexion plus inversion (dark blue), and inversion (light blue). The bars represent acceleration changes normalized by gravitational acceleration and averaged across subjects; error bars represent standard deviations across subjects. The left column represents changes using the muscle-driven models, and the right column represents changes using the torque-driven models. Asterisks above bars represent statistically significant changes relative to normal walking condition; diamonds above bars in the right column represent changes from torque-driven simulations that were statistically different from changes from muscle-driven simulations. The maximum acceleration change observed across both muscle-driven and torque-driven conditions was 0.93 m s^-2^.

**Fig S7.**
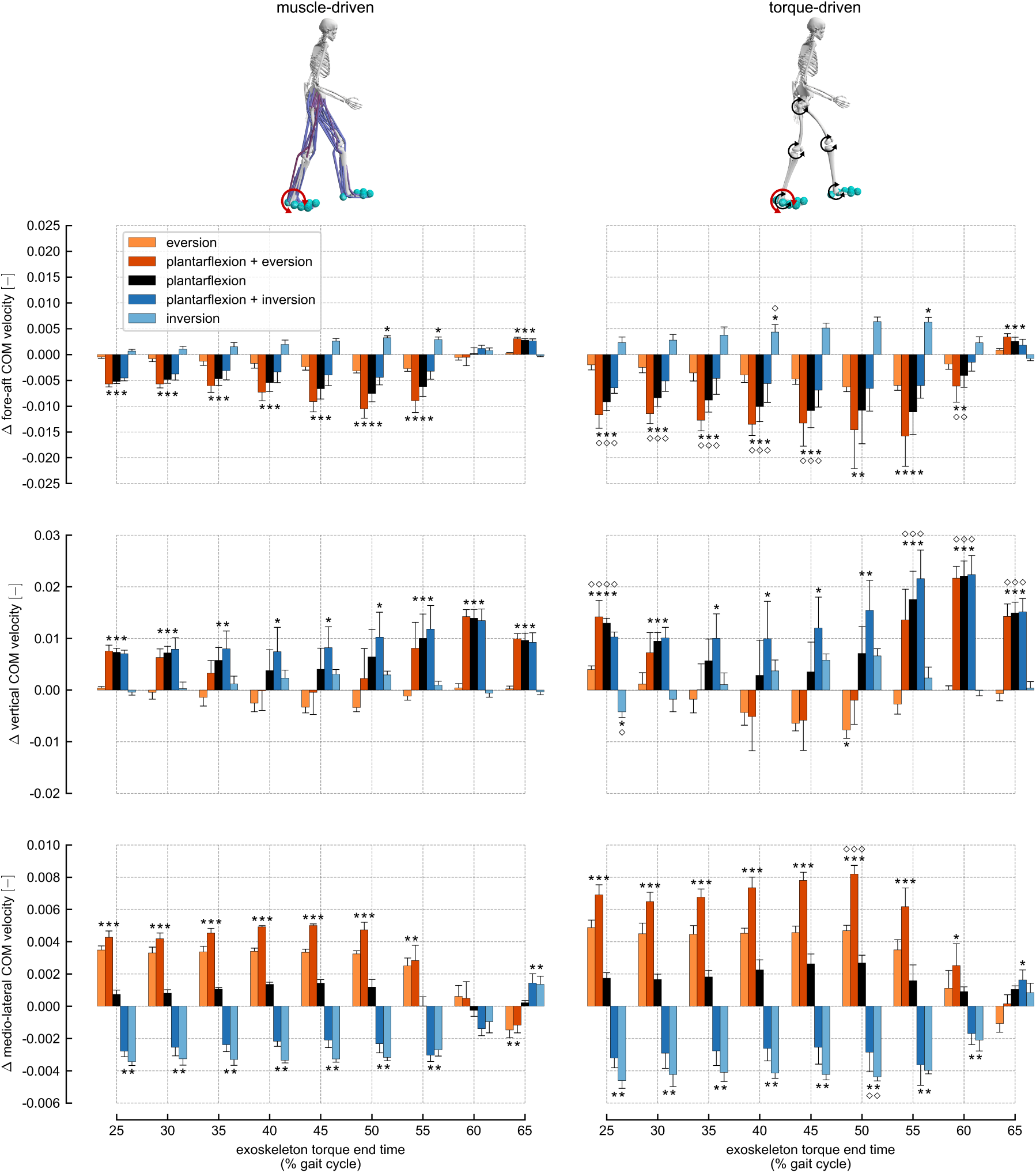
Changes in center of mass velocity. The change in center of mass velocity, calculated at exoskeleton torque end time, for each exoskeleton torque condition: eversion (light orange), plantarflexion plus eversion (dark orange), plantarflexion (black), plantarflexion plus inversion (dark blue), and inversion (light blue). Velocity changes are normalized to the dimensionless Froude number. The bars represent normalized velocity changes averaged across subjects; error bars represent standard deviations across subjects. The left column represents changes using the muscle-driven models, and the right column represents changes using the torque-driven models. Asterisks above bars represent statistically significant changes relative to normal walking condition; diamonds above bars in the right column represent changes from torque-driven simulations that were statistically different from changes from muscle-driven simulations. The maximum velocity change observed across both muscle-driven and torque-driven conditions was 0.072 m s^-1^.

**Fig S8.**
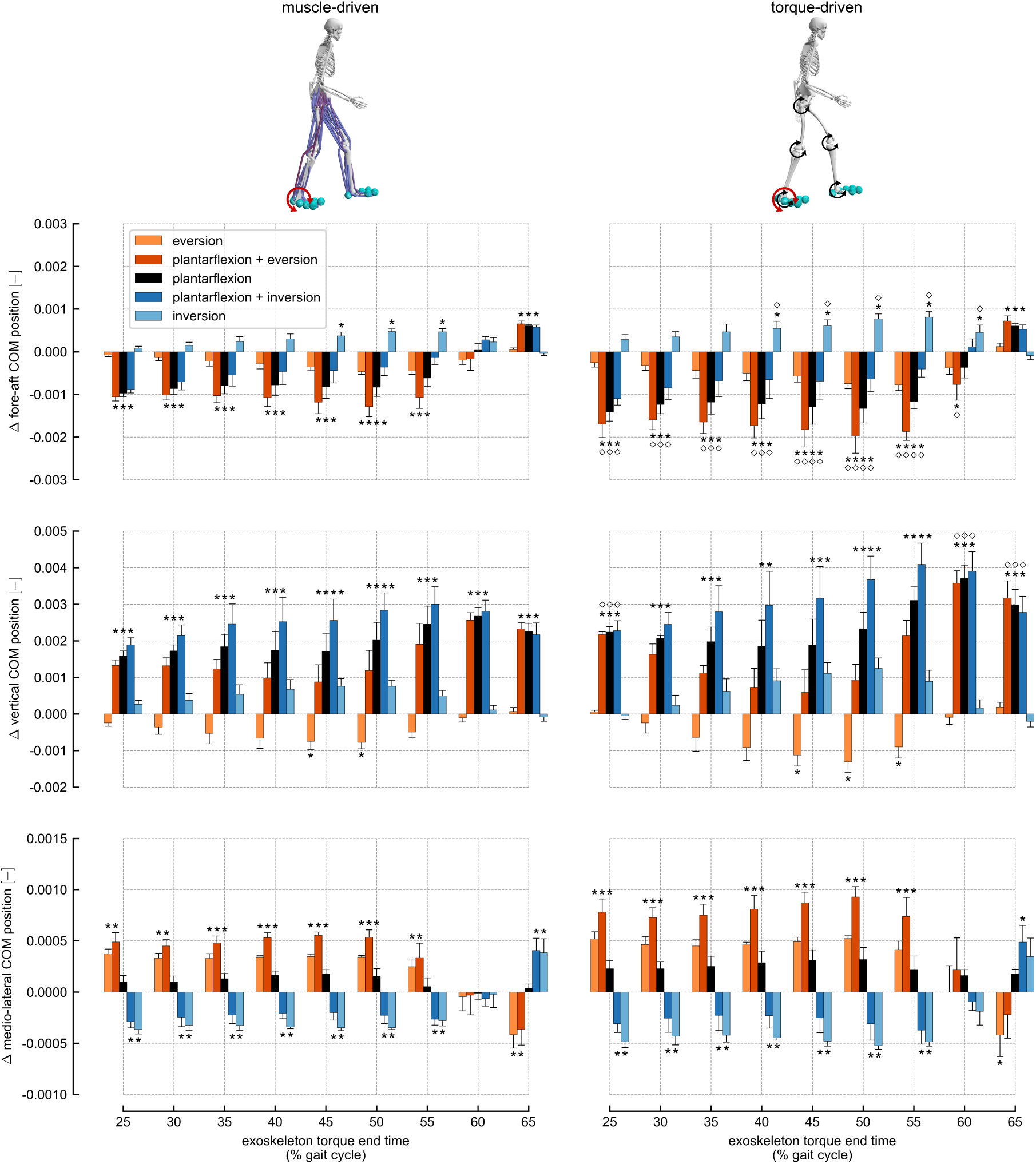
Changes in center of mass position. The change in center of mass position, calculated at exoskeleton torque end time, for each exoskeleton torque condition: eversion (light orange), plantarflexion plus eversion (dark orange), plantarflexion (black), plantarflexion plus inversion (dark blue), and inversion (light blue). The bars represent position changes averaged across subjects; error bars represent standard deviations across subjects. The left column represents changes using the muscle-driven models, and the right column represents changes using the torque-driven models. Asterisks above bars represent statistically significant changes relative to normal walking condition; diamonds above bars in the right column represent changes from torque-driven simulations that were statistically different from changes from muscle-driven simulations. The maximum position change observed across both muscle-driven and torque-driven conditions was 0.004 m.

**Fig S9.**
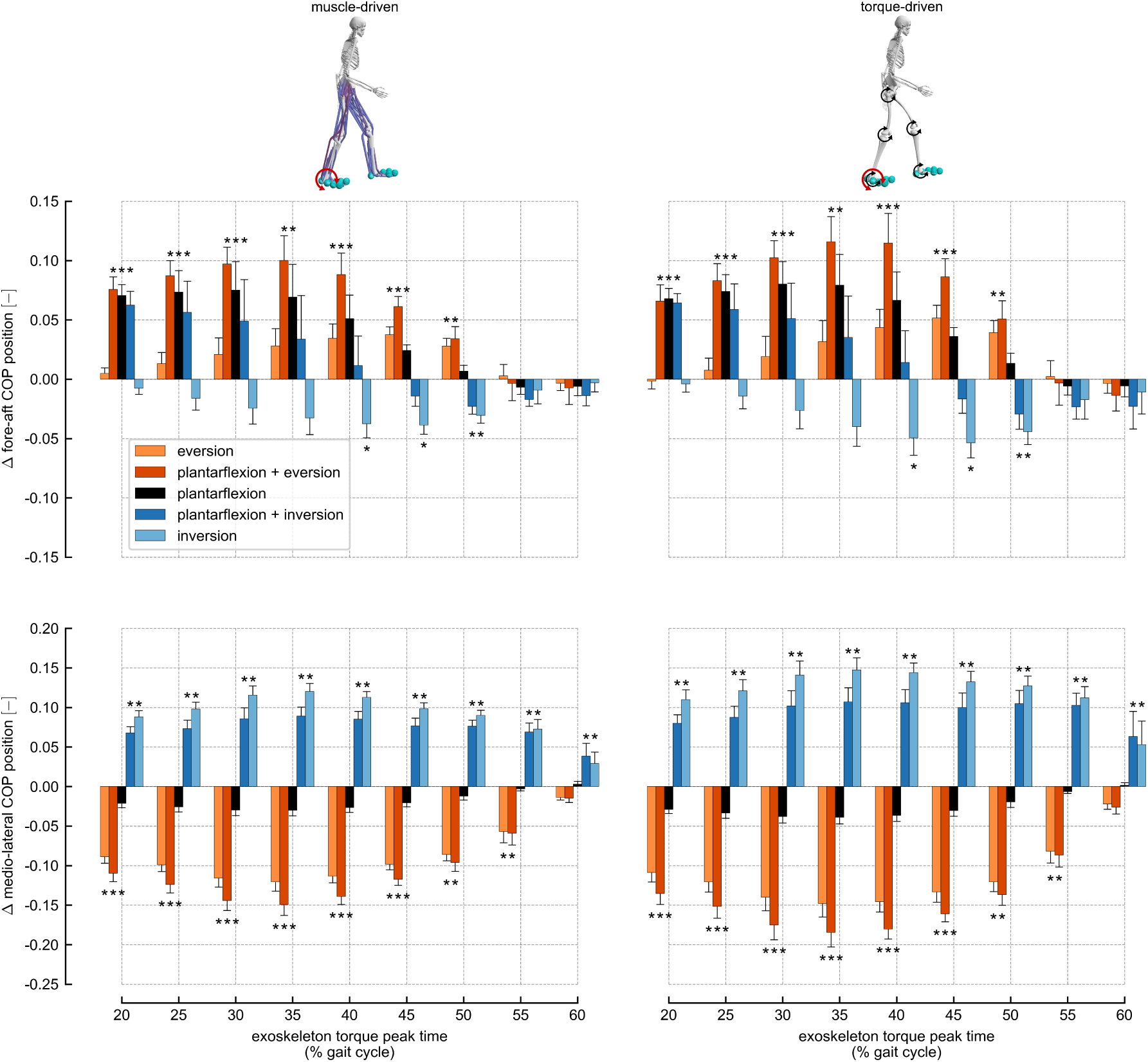
Changes in right foot center of pressure position. The change in right foot center of pressure position, calculated at the exoskeleton torque peak time, for each exoskeleton torque condition: eversion (light orange), plantarflexion plus eversion (dark orange), plantarflexion (black), plantarflexion plus inversion (dark blue), and inversion (light blue). The bars represent position changes averaged across subjects; error bars represent standard deviations across subjects. The left column represents changes using the muscle-driven models, and the right column represents changes using the torque-driven models. Asterisks above bars represent statistically significant changes relative to normal walking condition. No significant differences in center of pressure changes between torque-driven and muscle-driven simulations were detected. The maximum center of pressure position change observed across both muscle-driven and torque-driven conditions was 0.024 m.

**Fig S10.**
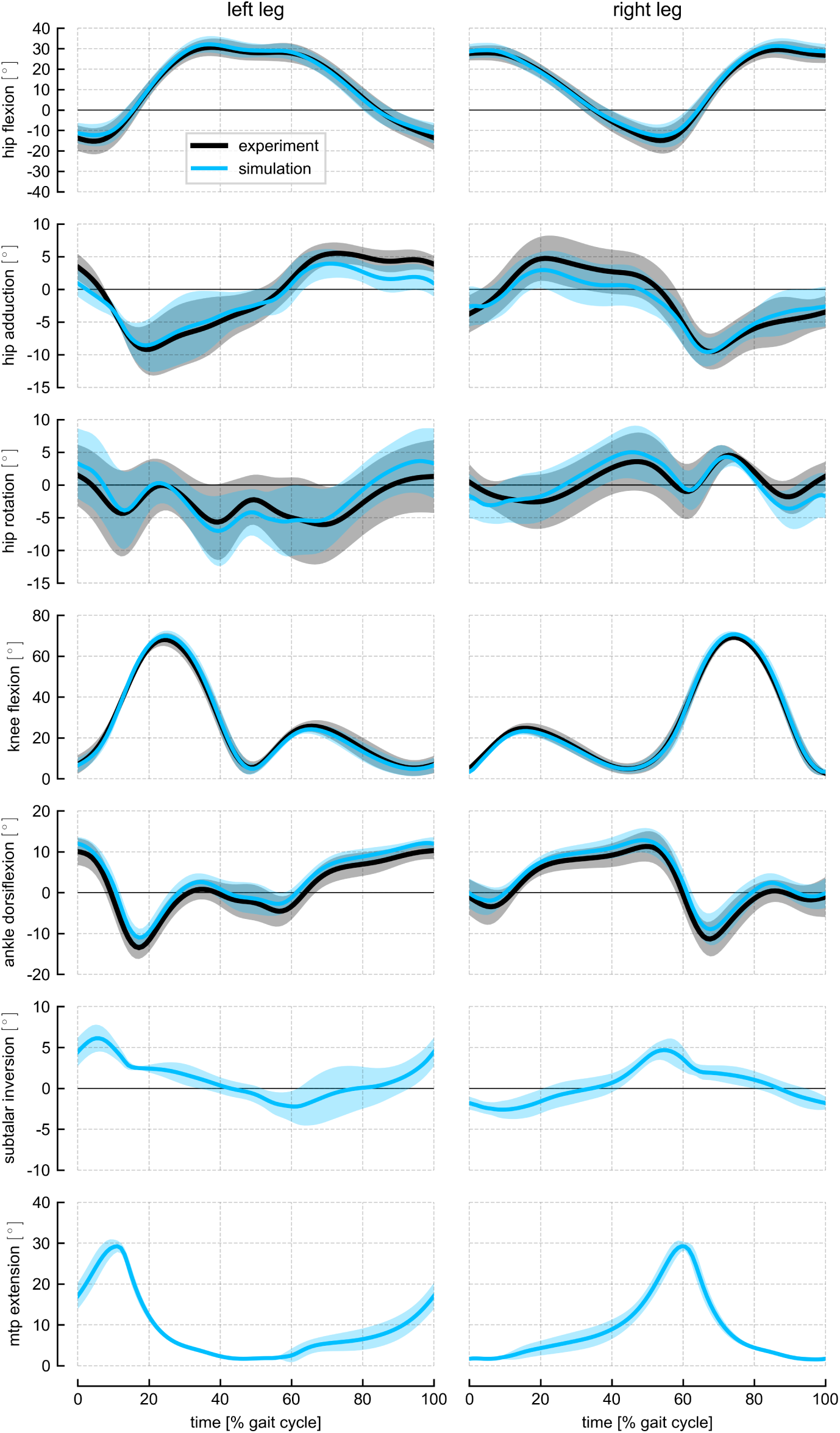
Lower-limb joint angles. Joint angles computed from experimental data using inverse kinematics (black) compared to joint angles from the tracking simulations (blue). Solid lines represent averages across subjects, and shaded bands represent standard deviations across subjects. The subtalar and metatarsophalangeal (mtp) joints did not track any reference experimental data.

**Fig S11.**
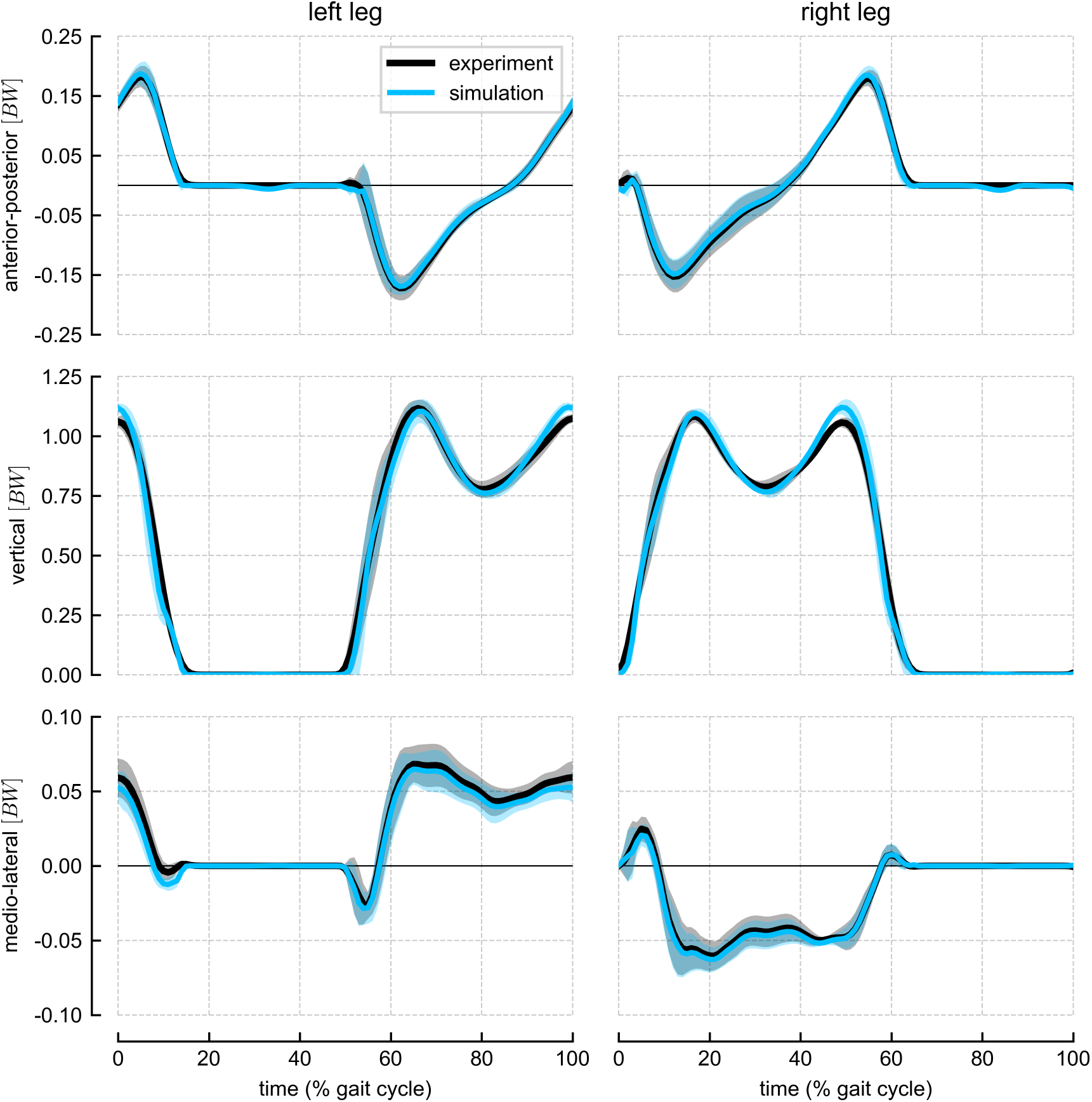
Ground reaction forces. Experimental ground reaction forces (black) compared to model-generated ground reaction forces from the tracking simulations (blue). Solid lines represent averages across subjects, and shaded bands represent standard deviations across subjects.

**Fig S12.**
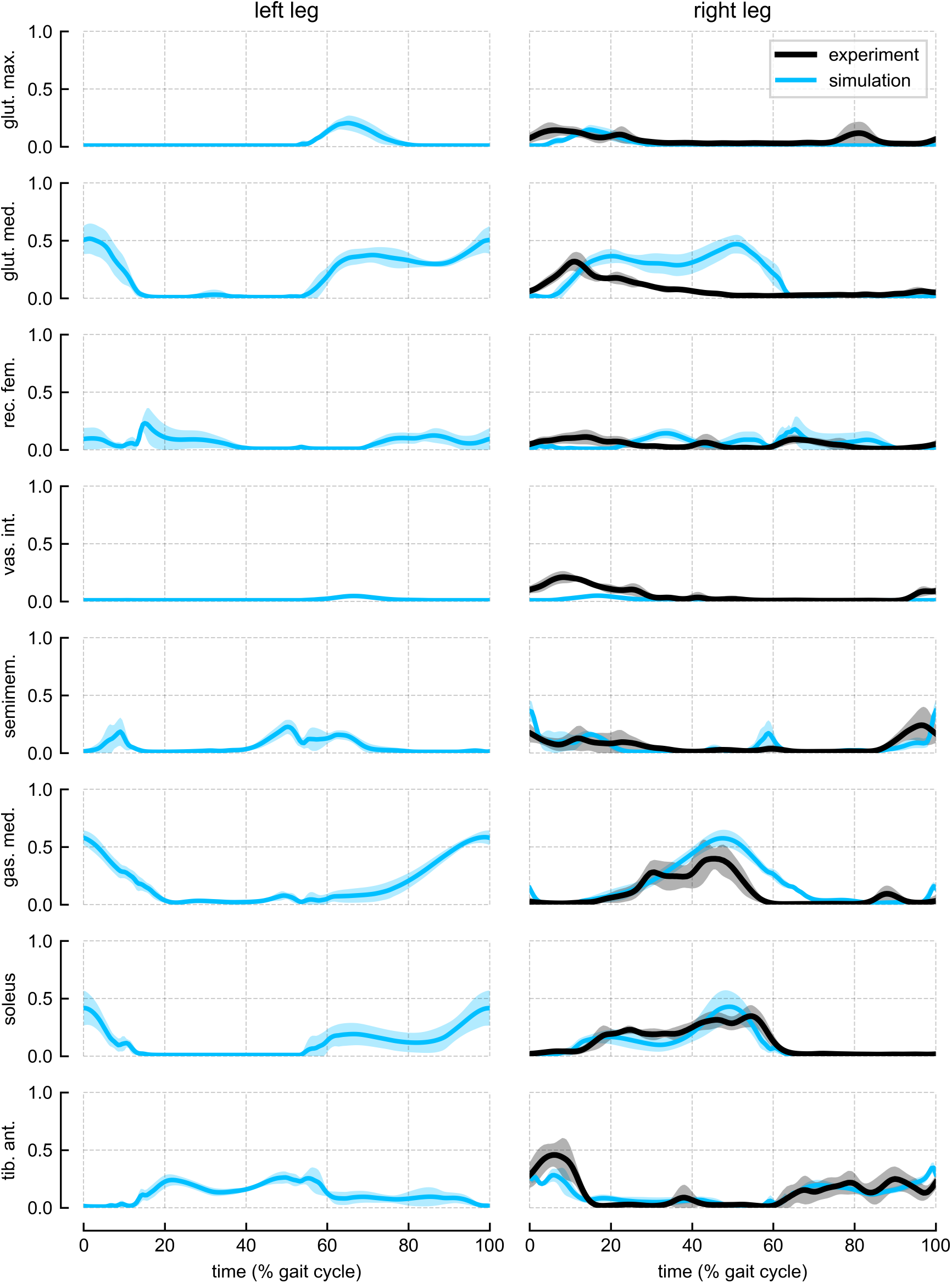
Muscle activations versus electromyography. Experimental electromyography recordings (black, right leg only) compared to muscle activations from the tracking simulations (blue). To account for electromechanical delays, a 40 ms delay was applied to the experimental EMG recordings. Solid lines represent averages across subjects, and shaded bands represent standard deviations across subjects.

**Fig S13.**
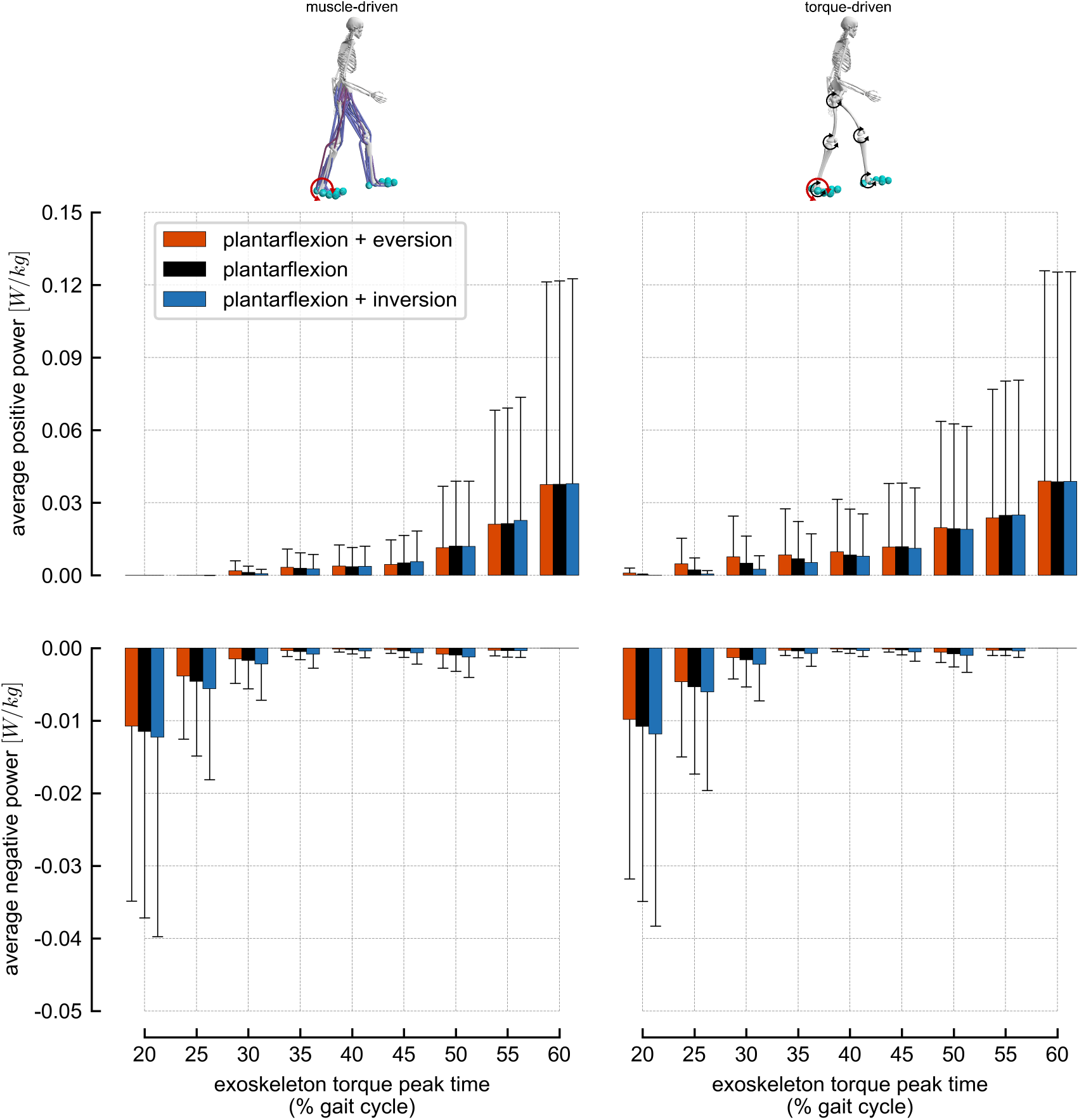
Plantarflexion exoskeleton powers. Average positive and average negative plantarflexion exoskeleton powers for the following exoskeleton torque conditions: plantarflexion plus eversion (dark orange), plantarflexion (black), plantarflexion plus inversion (dark blue). Power is averaged over the full simulation time range when exoskeleton torque was applied. Exoskeleton timings are denoted by the time of peak exoskeleton torque. For the plantarflexion plus eversion and plantarflexion plus inversion exoskeletons, only the power produced by the plantarflexion torque component is shown. Each bar represents the average positive (top) or average negative (bottom) power normalized by body mass and averaged across subjects; error bars represent standard deviations across subjects. The left column represents changes using the muscle-driven models, and the right column represents changes using the torque-driven models. Torques during push-off (60% of the gait cycle) produced the largest average positive powers, which corresponded with the largest forward center of mass velocity changes we observed. Early mid-stance torques (20% of the gait cycle) produced the largest average negative powers. Backward changes in center of mass velocity were observed throughout mid-stance, but large average negative powers did not persist beyond early mid-stance.

**Fig S14.**
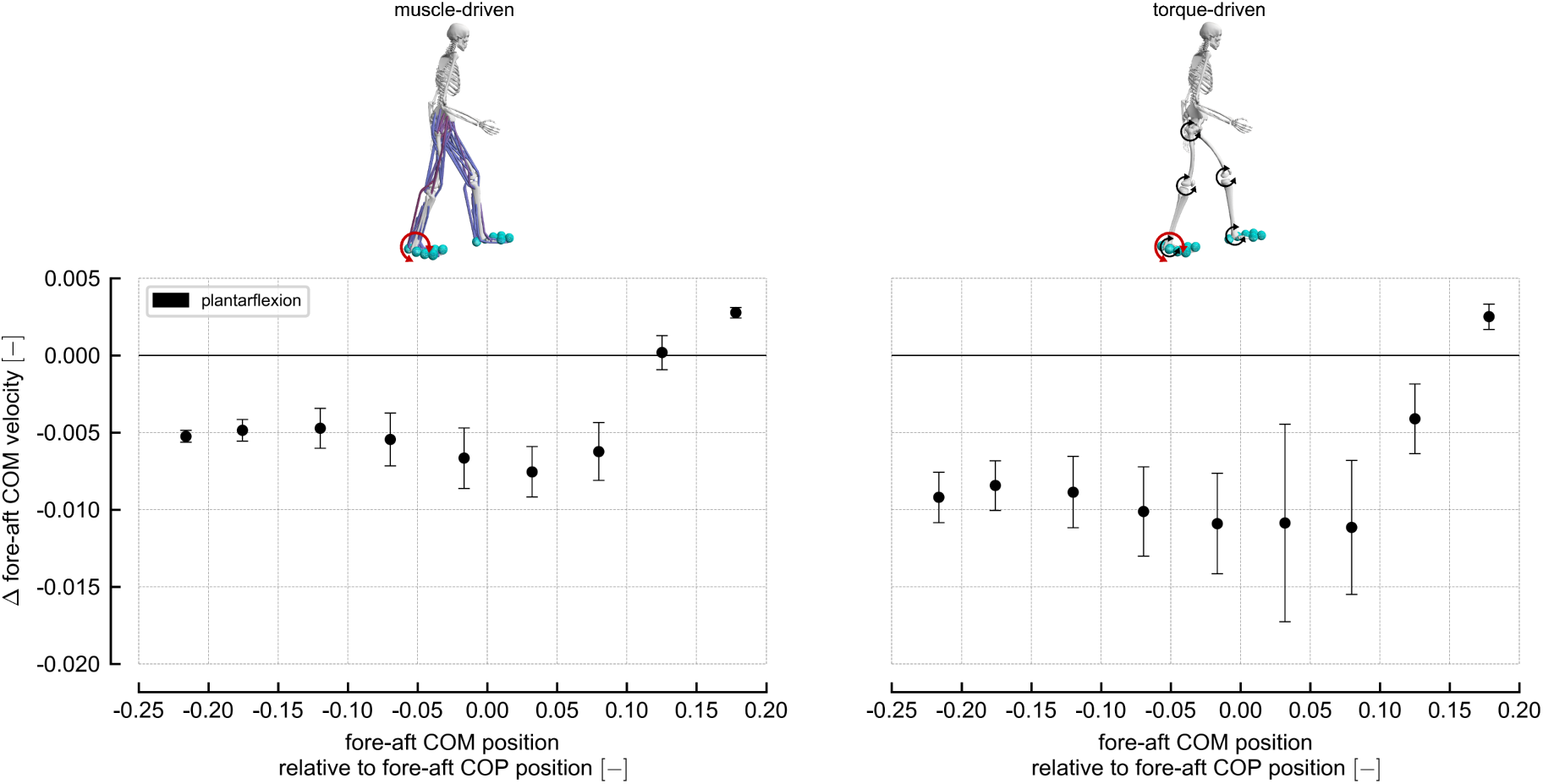
Fore-aft center of mass velocity changes versus fore-aft center of mass position. The change in fore-aft center of mass velocity, calculated at exoskeleton torque end time, for the plantarflexion exoskeleton torque condition. The x-axis represents the fore-aft center of mass position relative to the fore-aft center of pressure position, normalized by center of mass height and calculated at exoskeleton torque onset time. The filled circles represent mean values across subjects; error bars represent standard deviations across subjects. The left column represents changes using the muscle-driven models, and the right column represents changes using the torque-driven models. From left to right, the circles represent measurements from 25% to 65% of the gait cycle, corresponding to the nine exoskeleton timings we simulated. As the center of mass passed in front of the center of pressure (positive x-axis values), the velocity changes remained negative (backward) until the latest exoskeleton timings. This suggests that changes in fore-aft center of mass velocities were not strongly related to the position of the center of mass relative to the center of pressure.

**Fig S15.**
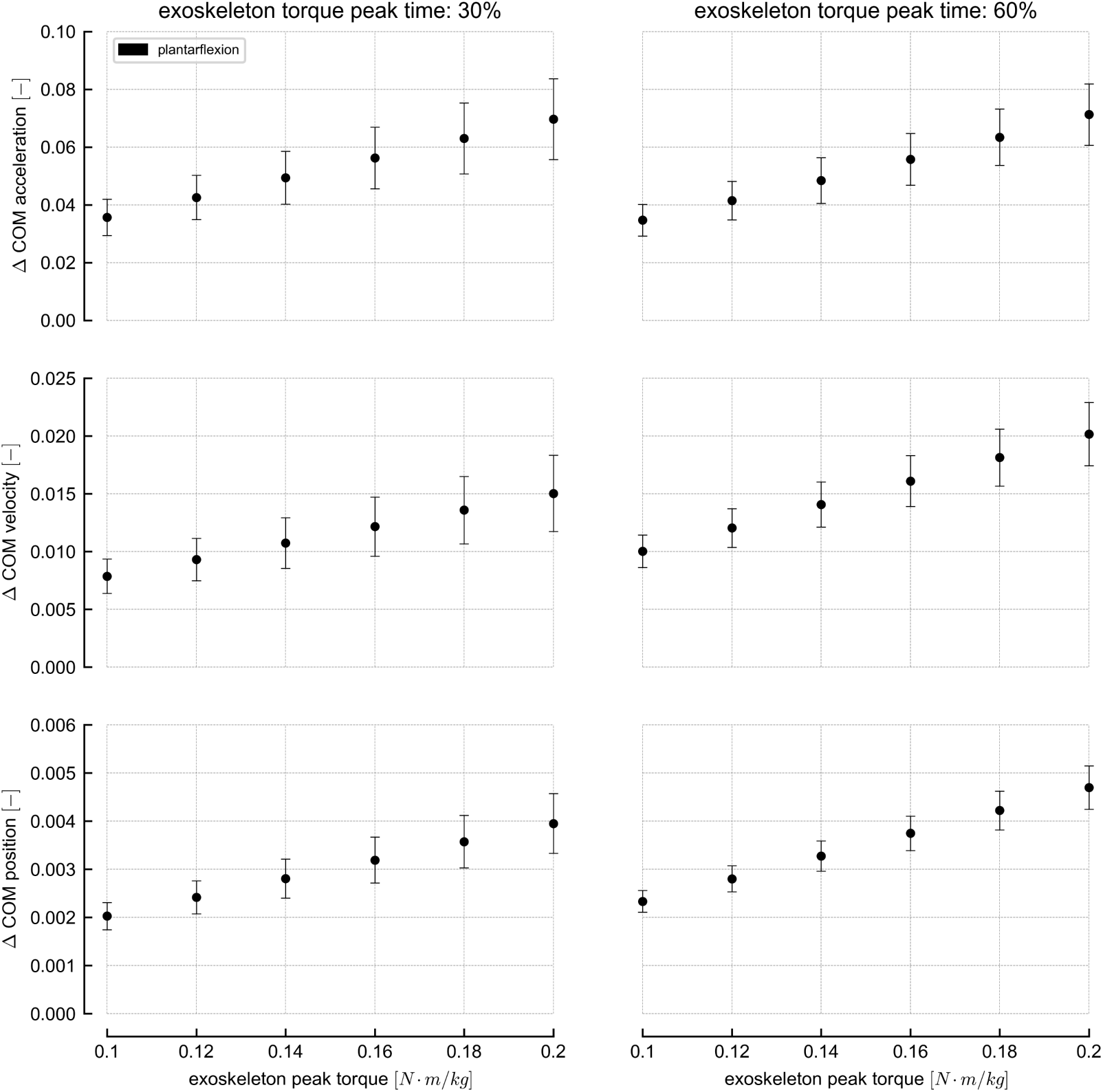
Center of mass changes versus exoskeleton peak torque. The changes in the 2-norm of the 3- dimensional center of mass position, velocity, and acceleration for the ankle plantarflexion exoskeleton torque condition versus exoskeleton peak torque. The kinematic changes were computed relative to the exoskeleton torque profile as described in Fig 1, and each quantity is normalized to be dimensionless based on the recommendations of Hof (1996). The filled circles represent mean values across subjects; error bars represent standard deviations across subjects. These results suggest that changes in these kinematic quantities have a strong linear relationship with exoskeleton peak torque.

**Fig S16.**
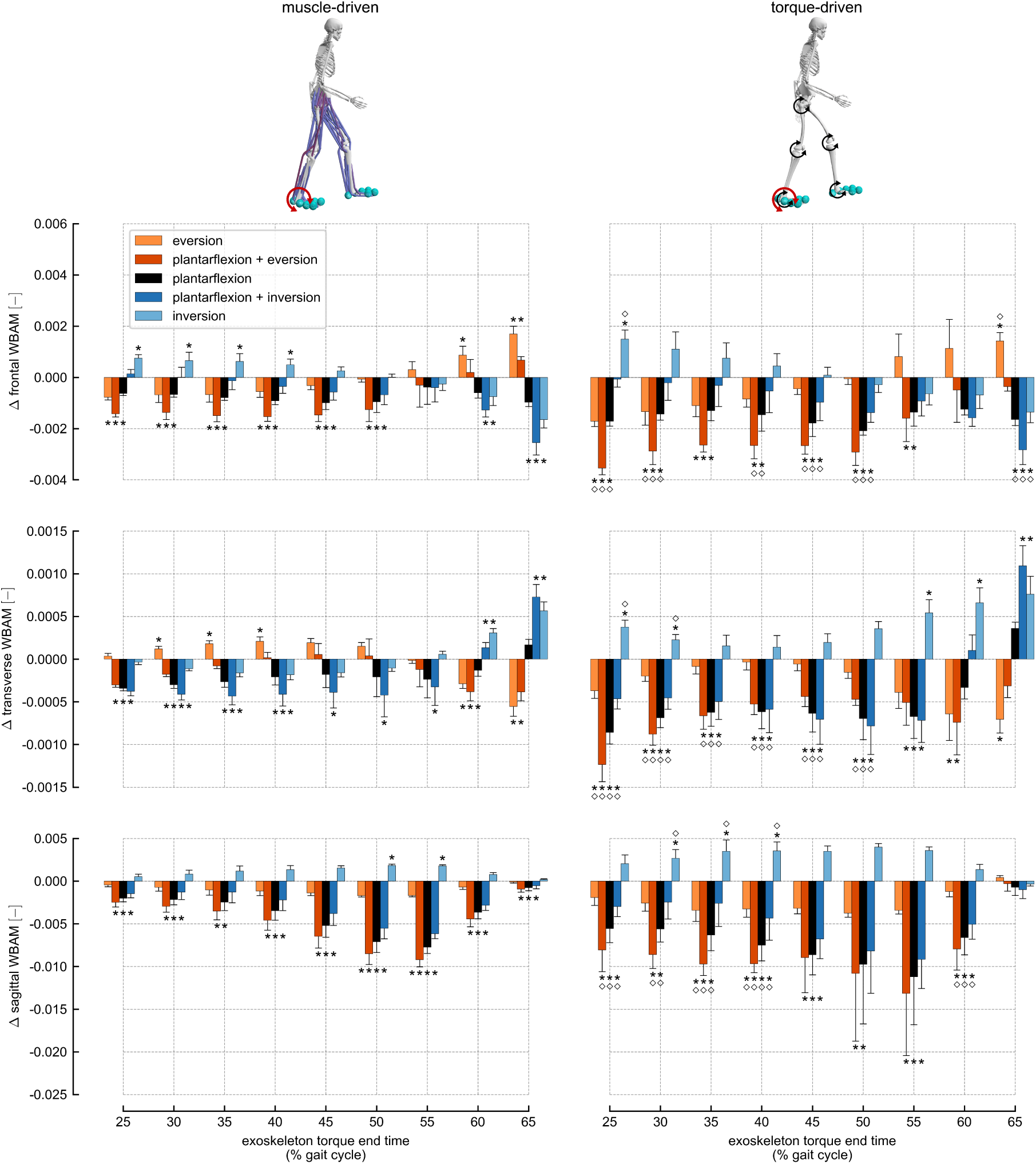
Changes in whole-body angular momentum. The change in whole-body angular momentum (WBAM), calculated at exoskeleton torque end time, for each exoskeleton torque condition: eversion (light orange), plantarflexion plus eversion (dark orange), plantarflexion (black), plantarflexion plus inversion (dark blue), and inversion (light blue). WBAM was normalized by mass, center of mass height, and 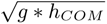 [76, 77]. The bars represent WBAM changes averaged across subjects; error bars represent standard deviations across subjects. The left column represents changes using the muscle-driven models, and the right column represents changes using the torque-driven models. Asterisks above bars represent statistically significant changes relative to normal walking condition; diamonds above bars in the right column represent changes from torque-driven simulations that were statistically different from changes from muscle-driven simulations. The maximum WBAM change observed across both muscle-driven and torque-driven conditions was 3.25 kg m2 s-1.

## Notes

### Competing Interest Statement

The authors have declared no competing interest.

### Summary of Updates

Various updates based on journal review responses. Add supplementary Figures S15 and S16. Revisions to clarify scientific impact and more thoroughly address previous exoskeleton work on balance.

